# Defining a Biofabrication Window for Visible-Light Photocrosslinkable Protein Bioinks in Extrusion Bioprinting

**DOI:** 10.64898/2026.09.15.751762

**Authors:** Jorge Martin-Perez, Sofia Martinez-Rodriguez, Cristina Castro-Dominguez, Adrian Sanz- Somoza, Alonso del Pozo-Dominguez, Gianna Arencibia, Fivos Panetsos

## Abstract

Extrusion bioprinting requires simultaneous control of material processability, rapid post-deposition stabilization, geometric fidelity, and cell compatibility. Here, we defined an experimental biofabrication window for three visible-light photocrosslinkable protein formulations based on silk fibroin (SF), gelatin methacryloyl (GelMA), and collagen methacryloyl (ColMA) using a extrusion-bioprinting workflow and ruthenium/sodium persulfate (Ru/SPS) photochemistry. Material-specific photocrosslinking conditions were first established under 430-nm irradiation and subsequently evaluated by ATR-FTIR spectroscopy, unconfined compression, short-term ARPE-19 cytocompatibility, and printing assays. Stable bulk hydrogels were obtained using 1× Ru/SPS, 45 mW, and 5 min for SF; 2.5× Ru/SPS, 45 mW, and 7 min for GelMA; and 2.5× Ru/SPS, 45 mW, and 6 min for ColMA. The selected formulations formed compliant hydrogels with initial compressive moduli of 0.60 ± 0.14, 0.52 ± 0.09, and 2.55 ± 2.20 kPa for SF, GelMA, and ColMA, respectively, without significant differences among groups. Short-term XTT assays identified the unreacted Ru/SPS-containing precursor as a major biological constraint, whereas photoinitiator-free SF and GelMA maintained metabolic activity close to the culture-medium reference. GelMA and SF generated interconnected lattices with printability indices of 0.893 ± 0.032 and 0.963 ± 0.081, respectively. Detailed SF analysis showed high positional accuracy (96.9 ± 4.8%) despite substantial post-deposition spreading, demonstrating that positioning accuracy and dimensional fidelity are distinct properties. Cell-laden printing was achieved with both SF and GelMA, and independent extrusion testing showed no statistically significant reduction in short-term ARPE-19 viability after SF bioprinting. Following the complete printing and photocrosslinking workflow, both formulations contained predominantly viable cells, but only SF retained the printed grid-associated cellular architecture under the tested conditions. These findings support the concept that biofabrication performance is an operational property arising from the intersection of photocrosslinking, cytocompatibility, extrusion compatibility, geometric fidelity, and post-fabrication structural stability.

## 1. Introduction

Tissue engineering seeks to restore, maintain, or replace damaged biological tissues by combining cells, biomaterials, and bioactive molecules within engineered microenvironments [1]. Conventional scaffold-based approaches have provided important advances in regenerative medicine, but they offer limited control over the spatial organization of cells and extracellular matrix-like materials. Three-dimensional bioprinting addresses this limitation by enabling the automated deposition of biomaterials and living cells according to predefined digital geometries, thereby providing control over construct architecture, cellular distribution, and material composition [2,3]. Among the available biofabrication strategies, extrusion-based bioprinting has become one of the most widely adopted because it is compatible with a broad range of hydrogel formulations, can process relatively high cell densities, and enables continuous deposition of material into predefined patterns [2,4].

The versatility of extrusion bioprinting is accompanied by a fundamental process constraint: successful deposition requires simultaneous control of material flow through the nozzle and preservation of the deposited geometry after extrusion. Printability therefore cannot be described solely by whether a formulation can pass through a nozzle. It encompasses extrudability, printing accuracy, and shape fidelity, all of which depend on both the physicochemical properties of the bioink and the operating conditions of the printing process [4,5]. Parameters including viscosity, shear-thinning behaviour, yield stress, recovery after shear, extrusion rate, printing speed, and nozzle diameter can determine whether deposition results in continuous filament formation, discontinuous flow, clogging, excessive spreading, or structural collapse [5–7]. These factors are also biologically relevant because the shear stresses generated during extrusion can compromise cell integrity, particularly as nozzle diameter decreases or flow resistance increases [8]. Consequently, extrusion-based biofabrication involves an intrinsic trade-off between processability, geometric resolution, structural stability, and cell compatibility.

This trade-off is particularly important for hydrogel bioinks. Their high water content and extracellular matrix-like characteristics make hydrogels attractive matrices for cell encapsulation and tissue engineering, but many biologically favourable formulations are mechanically weak before crosslinking and rapidly spread after deposition [4,5]. Gelatin methacryloyl (GelMA), collagen methacryloyl (ColMA), and silk fibroin represent three protein-based systems with distinct molecular structures and gelation mechanisms that are relevant in this context. GelMA is obtained by functionalizing gelatin with methacryloyl groups, combining cell-interactive motifs inherited from collagen with photocrosslinkable functionality. Its mechanical properties and network density can be modulated through polymer concentration, degree of functionalization, and crosslinking conditions, making it one of the most extensively studied photocrosslinkable bioinks [9]. Type I collagen provides a highly bioactive extracellular matrix-derived environment but forms comparatively weak hydrogels and presents challenges associated with controlled gelation and structural stability. Methacrylation enables formation of ColMA networks through radical-mediated photocrosslinking and improves control over hydrogel stabilization [10]. ColMA has consequently been investigated as a printable matrix for cell-containing tissue-engineered constructs [11].

Silk fibroin provides a complementary protein platform with substantially different molecular behaviour. Regenerated *Bombyx mori* silk fibroin can be processed in aqueous solution and subsequently stabilized through covalent and conformational mechanisms while retaining favourable biocompatibility, tunable mechanical properties, and controllable degradation [12]. Silk-based formulations have previously been incorporated into cell-laden bioprinting systems and have supported the survival and differentiation of encapsulated cells [12]. However, the behaviour of regenerated silk fibroin depends strongly on processing history, protein concentration, conformational state, sterilization, ionic environment, and gelation conditions. These variables can influence not only hydrogel formation but also extrusion behaviour and structural stabilization after deposition. More generally, accurate execution of a programmed toolpath does not necessarily guarantee preservation of the intended final geometry, because spreading and deformation may occur between material deposition and complete network stabilization. Thus, positional deposition accuracy and post-print shape fidelity should be regarded as related but distinct properties of an extrusion-biofabrication process.

Rapid post-deposition stabilization is therefore a central requirement when working with soft or low-viscosity protein bioinks. Photocrosslinking provides temporal and spatial control over hydrogel formation and can limit material spreading by converting the extruded precursor into a crosslinked network shortly after deposition [5,13]. Visible-light photoinitiating systems are particularly attractive for cell-containing formulations because they avoid reliance on shorter-wavelength ultraviolet irradiation and can provide greater light penetration under appropriate conditions [14]. Among these systems, ruthenium/sodium persulfate (Ru/SPS) photochemistry has been used for the visible-light crosslinking of gelatin-, collagen-, and silk-based hydrogels [15–18]. Under visible-light excitation, the ruthenium complex enters a photoexcited state and participates in electron transfer with persulfate, generating reactive species capable of initiating covalent network formation. In methacrylated proteins such as GelMA and ColMA, these radicals initiate polymerization of methacryloyl double bonds, producing covalent crosslinks between polymer chains [15,18]. In native silk fibroin, by contrast, ruthenium-mediated oxidation of tyrosine residues promotes formation of tyrosyl radicals and subsequent dityrosine crosslinks between fibroin chains [16,17]. Silk stabilization may additionally evolve through protein self-assembly and β-sheet formation, meaning that the final network can reflect an interplay between rapid covalent crosslinking and slower conformational rearrangement [17].

The use of a common photocrosslinking chemistry does not, however, imply that chemically distinct protein formulations can be processed under identical conditions. Polymer concentration, reactive-group density, photoinitiator concentration, irradiation dose, extrusion behaviour, and post-deposition spreading collectively determine whether a formulation can be converted into a stable printed construct. These variables can also influence cytocompatibility. In particular, conditions that favour rapid or extensive network formation may simultaneously increase cellular exposure to initiating components before photocrosslinking is completed. The practically useful operating space of a cell-containing photocrosslinkable formulation is therefore bounded by several interdependent requirements. Here, we define this experimentally accessible combination of formulation, extrusion, stabilization, structural, and biological conditions as the biofabrication window.

Direct comparison of bioinks reported in separate studies is complicated by differences in polymer concentration, printing hardware, nozzle geometry, crosslinking chemistry, irradiation conditions, biological models, and definitions of printability [4–6]. Evaluating distinct protein formulations within a shared experimental platform can instead provide a controlled framework for identifying where formulation-specific and process-related constraints arise during successive stages of biofabrication. In the present study, we used a proprietary in-house extrusion-bioprinting platform to investigate visible-light photocrosslinkable formulations based on silk fibroin, GelMA, and ColMA. Rather than attempting to establish an intrinsic ranking among the three biomaterials, we examined the conditions required for each formulation to progress through a common biofabrication workflow. All three protein systems were evaluated for Ru/SPS-mediated photocrosslinking, physicochemical characteristics, compressive mechanical behaviour, and short-term cytocompatibility. GelMA and silk fibroin were subsequently taken forward for quantitative printability assessment and proof-of-concept cell-laden extrusion, while independent experiments were used to examine geometric deposition fidelity and cell viability associated with extrusion and the complete photocrosslinked biofabrication process. By integrating these endpoints, the study aimed to identify the interacting material, processing, structural, and biological constraints that define a usable biofabrication window for visible-light photocrosslinkable protein bioinks.

## 2. Materials and Methods

### 2.1 Study design

This study was designed to define the experimental biofabrication window of three protein-based formulations processed within a common extrusion-bioprinting workflow: gelatin methacryloyl (GelMA), collagen methacryloyl (ColMA), and regenerated *Bombyx mori* silk fibroin (SF). All three systems were evaluated using a ruthenium/sodium persulfate (Ru/SPS) visible-light initiating system and irradiation at 430 nm. The main experimental work was conducted at Silk Biomed S.L. (Madrid, Spain).

The experimental workflow was organized sequentially (Figure 1). First, the extrusion system and the material-delivery configuration were established. The protein formulations were then screened to identify material-specific combinations of Ru/SPS concentration, optical power, and irradiation time capable of producing macroscopically stable hydrogels. The selected formulations were characterized by attenuated total reflectance Fourier-transform infrared spectroscopy (ATR-FTIR) and unconfined uniaxial compression. Cytocompatibility was subsequently evaluated in ARPE-19 cells by metabolic activity and fluorescence-based morphological assessment. GelMA and SF were then assessed using a standardized lattice-printing assay. A separate SF dataset was used to distinguish positional deposition accuracy from post-deposition geometric deformation. Finally, cell-containing formulations were evaluated through spatial cell localization, an independent extrusion-specific Live/Dead assay, and same-day viability imaging after the complete Ru/SPS-mediated printing and photocrosslinking process.

**Figure 1.**
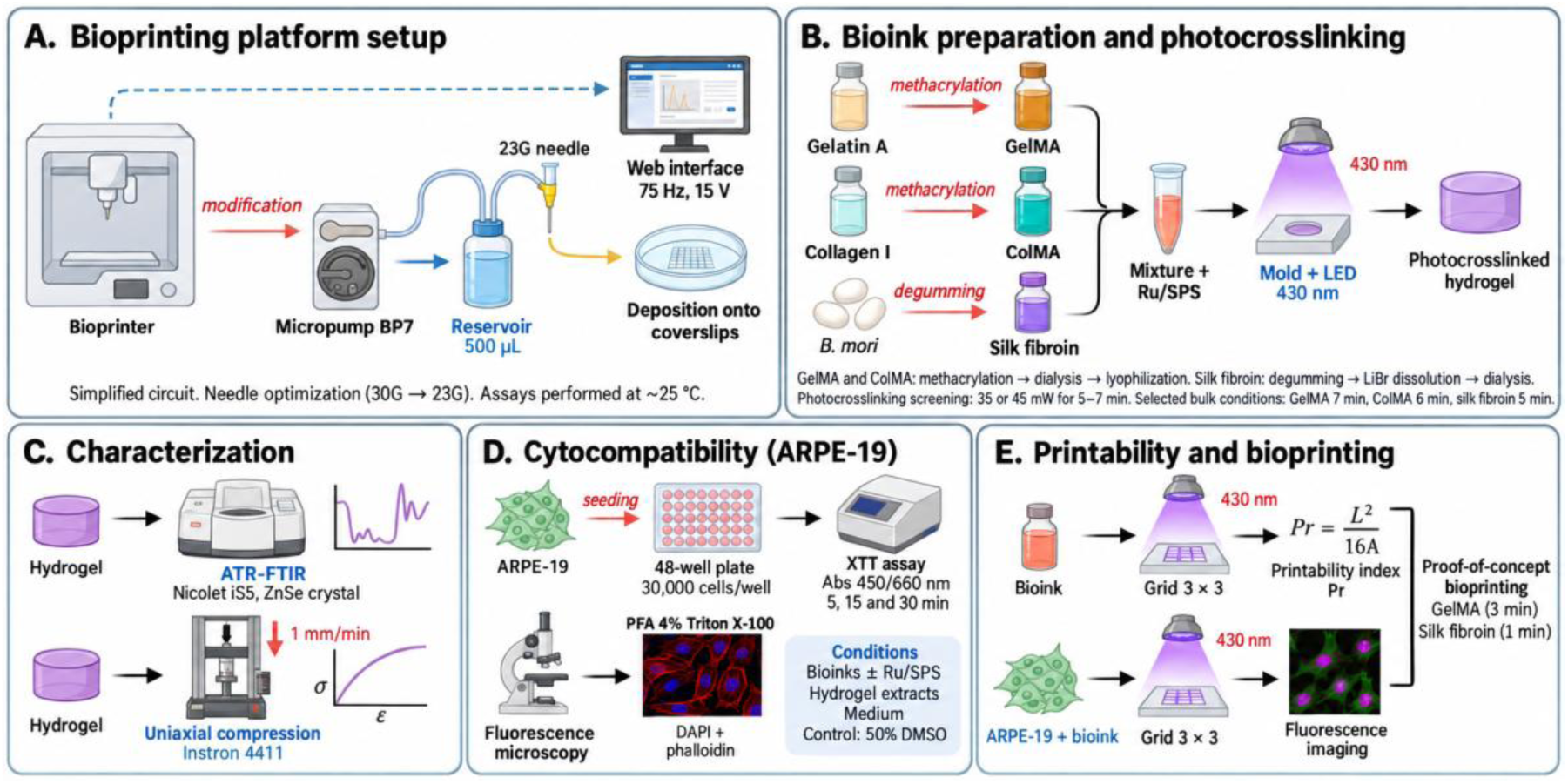
Overview of the experimental workflow. (A) Optimization of the in-house extrusion-bioprinting system. The initial fluidic configuration was simplified by removing the microfluidic chip and replacing the conventional 30G outlet needle with a 23G blunt needle. The final fluidic path comprised the micropump, connecting tubing, bioink reservoir, adapter, downstream tubing, and extrusion needle. The system was controlled through a web-based interface using operating settings of approximately 75 Hz and 15 V, and printing experiments were performed at approximately 25 °C. (B) Preparation and visible-light photocrosslinking of the protein formulations. Type-A gelatin and type-I collagen were methacrylated to obtain GelMA and ColMA, respectively, whereas SF was regenerated from Bombyx mori cocoons. The formulations were combined with Ru/SPS and screened under 430-nm irradiation. (C) Physicochemical and mechanical characterization by ATR-FTIR spectroscopy and unconfined uniaxial compression. (D) Cytocompatibility assessment in ARPE-19 cells by XTT metabolic activity and fluorescence microscopy following DAPI and phalloidin staining. (E) Biofabrication assessment comprising 3 × 3 lattice printing, printability-index analysis, quantitative geometric-fidelity analysis, cell-containing extrusion, and post-print fluorescence-based cell assessment. Created with BioRender.

ColMA was included in the photocrosslinking, ATR-FTIR, mechanical, and cytocompatibility experiments. Because of the limited quantity of material recovered after synthesis, it was not included in the final lattice-printing or cell-laden printing experiments.

### 2.2. Extrusion-bioprinting platform

Bioprinting experiments were performed using the THOR-X Bioprinter, a proprietary in-house extrusion platform developed by Omnia Mater S.L. The system incorporates computer-controlled positioning and a microfluidic pumping unit whose actuation frequency and voltage can be independently adjusted. Engineering features unrelated to reproduction of biomaterial experiments are subject to intellectual-property protection and are therefore not described in detail.

The fluidic delivery configuration was optimized iteratively to obtain continuous material deposition while reducing recurrent obstruction and excessive dead volume. In the final configuration used for the integrated GelMA and SF experiments, the microfluidic chip present in earlier versions of the system was removed from the extrusion pathway, and the dispensing outlet was changed from a conventional 30G needle to a sterile 23G blunt needle. The final fluidic pathway consisted of a micropump, connecting tubing, bioink reservoir, adapter, downstream tubing, and the 23G extrusion needle. This configuration was used for the principal lattice-printing and cell-containing experiments.

The platform was controlled through a web-based interface. For the final lattice-printing workflow, the experimental records indicate actuation settings of approximately 75 Hz and 15 V. These settings were maintained constant within each experimental series. Printing was performed at room temperature, approximately 25 °C. Digital constructs were designed using Tinkercad (Autodesk) and converted into printable toolpaths using EasyWare (EasyThreed, v. K10) and Cura (UltiMaker, v. 3.6.0).

### 2.3. Preparation of protein formulations

#### 2.3.1. Gelatin methacryloyl

GelMA was synthesized from type-A gelatin from porcine skin (Sigma-Aldrich, Cat. No. G2500) using a procedure previously established in the laboratory and adapted from Pérez-Cortez *et al.* [19]. Gelatin was dissolved in phosphate-buffered saline (PBS) at 30 °C to obtain a 10% (w/v) solution. In the preparation used for the present study, 2 g gelatin was dissolved in 20 mL PBS. Methacrylic anhydride (Sigma-Aldrich, Cat. No. 276685) was subsequently added dropwise under continuous mixing (2 mL per 20 mL gelatin solution). The reaction was terminated by dilution with PBS.

The resulting solution was dialyzed against distilled water using 12-14 kDa molecular-weight-cutoff dialysis tubing (BioDesign Dialysis Tubing™ D306, Thermo Fisher Scientific, Cat. No. 12797486) for 5 days with daily changes. After dialysis, the material was frozen at −80 °C and lyophilized for 2 days using a LyoQuest freeze dryer (Telstar).

For the experiments reported here, the lyophilized GelMA was dissolved at 25-30 °C in complete DMEM/F12 culture medium to a final polymer concentration of 16.67 mg/mL. The degree of methacryloyl functionalization of the GelMA batch was not independently quantified and was therefore not included as an experimental variable.

#### 2.3.2. Collagen methacryloyl

ColMA was prepared from rat-tail type-I collagen (Sigma-Aldrich, Cat. No. C7661) by adapting the preparation procedure associated with the PhotoCol®-RUT system (Advanced BioMatrix, Cat. No. CC322). Type-I collagen was dissolved in acetic acid to obtain a 0.5% (w/v) collagen solution. Methacrylic anhydride was added dropwise under continuous agitation at an MA:collagen-solution ratio of 1:1000 (v/v), and the reaction mixture was stirred at 500 rpm for 24 h at room temperature.

Following methacrylation, the solution was dialyzed against distilled water for 3 days using 12-14 kDa molecular-weight-cutoff dialysis tubing (BioDesign Dialysis Tubing™ D306, Thermo Fisher Scientific, Cat. No. 12797486). The purified material was frozen at −80 °C for approximately 24 h and subsequently lyophilized for 24 h using a LyoQuest freeze dryer (Telstar). Because the resulting lyophilizate formed a compact mass, it was mechanically fragmented using sterile forceps before weighing.

For formulation, ColMA was initially solubilized at 5 mg/mL in 20 mM acetic acid at approximately 4 °C. The stock was then combined with neutralization solution (Advanced BioMatrix, Cat. No. CC322) and complete culture medium to obtain a final polymer concentration of 3 mg/mL. The material was maintained at approximately 4 °C during preparation to limit premature physical gelation. The degree of collagen methacrylation was not independently quantified.

#### 2.3.3. Silk fibroin

Regenerated aqueous SF was prepared from *Bombyx mori* cocoons, following a previously established procedure [20]. Cocoons were cut into approximately 2-3 cm² fragments and degummed in 0.2% (w/v) sodium carbonate (Na₂CO₃) for 20 min at 121 °C and 103.4 kPa. Following degumming, the fibers were extensively washed with distilled water, dried, and dissolved in 9.3 M lithium bromide (LiBr) for 3 h at 60 °C.

The resulting SF solution was dialyzed against distilled water for 3 days at 4 °C using 3.5 kDa molecular-weight-cutoff dialysis tubing (BioDesign Dialysis Tubing™, Thermo Fisher Scientific, Cat.

No. 12707496), with four to five water changes. Removal of residual salts was monitored by conductivity using a conductivity meter. Following dialysis, the solution was centrifuged at 4863 × *g* using a centrifuge to remove insoluble material. The recovered supernatant was concentrated by reverse dialysis to approximately 6.5% (w/v) and subsequently diluted in complete culture medium to a final concentration of 2% (w/v) for the main biofabrication experiments.

For experiments involving cells, all formulations were sterilized by filtration through 0.2-µm hydrophilic PES membranes (Millex™, Millipore, Cat. No. SLGPR33RS). This sterilization strategy was selected because previous optimization showed that filtration preserved visible-light-induced gelation, whereas autoclaving impaired subsequent photocrosslinking.

### 2.4. Culture medium and Ru/SPS photoinitiating system

Unless otherwise indicated, DMEM/F12 (Gibco, Cat. No. 11320-033) supplemented with 10% fetal bovine serum (FBS; Gibco, Cat. No. A5256701) and 1% penicillin/streptomycin (Gibco, Cat. No. 15140122) was used as complete culture medium. The same medium was used as the aqueous culture-medium component of the GelMA and SF formulations.

The ruthenium photoinitiator and sodium persulfate (SPS) were obtained from the PhotoCol®-RUT system (Advanced BioMatrix, Cat. No. CC322). Stock solutions were handled under reduced ambient-light exposure and added to the protein formulations shortly before irradiation. The reference Ru/SPS concentration, designated **1×**, corresponded to final concentrations of 0.31 mg/mL Ru and 0.99 mg/mL SPS. Higher conditions are expressed throughout the manuscript as proportional multiples of this reference concentration. The exact final composition of the selected formulations is reported in Table 1 in the Results section.

**Table 1.** Final protein formulations and visible-light photocrosslinking conditions selected for downstream bulk characterization. Polymer concentration, final ruthenium (Ru) and sodium persulfate (SPS) concentrations, formulation vehicle, optical power, and 430-nm irradiation time are shown for GelMA, ColMA, and silk fibroin (SF). The reference Ru/SPS condition, defined as 1×, corresponded to 0.31 mg/mL Ru and 0.99 mg/mL SPS; 2.5× corresponded to 0.78 mg/mL Ru and 2.48 mg/mL SPS..

| Bioink | Polymer concentration | Ru (mg/mL) | SPS (mg/mL) | Vehicle | Optical power | 430 nm irradiation |
| --- | --- | --- | --- | --- | --- | --- |
| GelMA | 16.67 mg/mL | 0.78 (2.5×) | 2.48 (2.5×) | DMEM/F12 medium | 45 mW | 7 min |
| ColMA | 3 mg/mL | 0.78 (2.5×) | 2.48 (2.5×) | Acetic acid / culture medium | 45 mW | 6 min |
| Silk fibroin | 2 % (w/v) | 0.31 (1×) | 0.99 (1×) | DMEM/F12 medium | 45 mW | 5 min |

### 2.5. Visible-light photocrosslinking and bulk formulation screening

Bulk photocrosslinking was evaluated using a mounted 430-nm LED source (M430L5, Thorlabs). Hydrogels were prepared by dispensing 200 µL of each formulation into open cylindrical molds with an internal diameter of 6 mm and a nominal height of 4.6 mm. The molds were produced by 3D printing (K1C 3D Printer, Creality, PLA as printing material) and sealed at the base with Parafilm. Irradiation was applied from above.

The illumination beam was focused onto an approximately 6-mm-diameter circular area encompassing the upper surface of the sample. Optical power at the sample position was verified using a PM16-121 optical power meter equipped with a silicon photodiode detector (Thorlabs). The final screening compared nominal optical powers of 35 and 45 mW and exposure times of 5, 6, and 7 min. Ru/SPS concentration was varied according to the protein formulation: SF was evaluated at 1× and 2×, with the higher 2.5× condition also assessed during formulation screening; GelMA was evaluated at 1×, 2×, and 2.5×; and the final ColMA condition was assessed at 2.5×. The material-specific conditions selected from this screening and subsequently used for downstream bulk characterization are reported in Table 1.

Photocrosslinking was considered operationally successful when a sample could be removed from the mold as an intact hydrogel without fragmentation and retained the imposed cylindrical geometry during manual handling. Hydrogels were recovered using a biopsy punch. This macroscopic endpoint was used solely as a practical criterion for selecting process conditions and was not interpreted as a quantitative measurement of crosslink density.

For the selected bulk formulations, the final irradiation conditions were 45 mW for 7 min for GelMA, 45 mW for 6 min for ColMA, and 45 mW for 5 min for SF. These conditions were used for the principal ATR-FTIR, mechanical, and cytocompatibility datasets.

### 2.6. ATR-FTIR spectroscopy

Chemical and conformational characteristics of the protein formulations were assessed by ATR-FTIR spectroscopy using a Nicolet iS5 FTIR spectrometer equipped with an iD5 ATR accessory and a zinc selenide crystal (Thermo Scientific). For each biomaterial, two states were compared: the uncrosslinked protein formulation without Ru/SPS and the corresponding formulation after addition of Ru/SPS and visible-light photocrosslinking under the selected bulk conditions.

Spectra were acquired in absorbance mode from 4000 to 600 cm⁻¹ at a spectral resolution of 4 cm⁻¹. A common background spectrum was acquired before sample measurements. Each individual spectrum was baseline-corrected by subtracting the mean absorbance measured between 1780 and 1800 cm⁻¹.

The complete baseline-corrected spectra were initially evaluated over the full acquisition range. For quantitative comparison of the protein-associated region, spectra between 1800 and 1200 cm⁻¹ were normalized to the Amide II band. Four spectral indices were subsequently calculated:

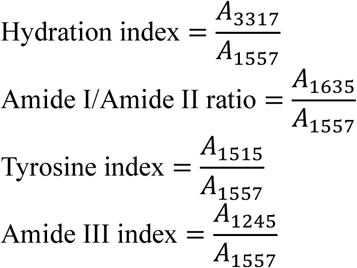

The final dataset comprised ColMA: n = 2 uncrosslinked and n = 2 photocrosslinked; GelMA: n = 3 uncrosslinked and n = 4 photocrosslinked; SF: n = 3 uncrosslinked and n = 3 photocrosslinked, based on the datasets represented in Figure 3.

**Figure 2.**
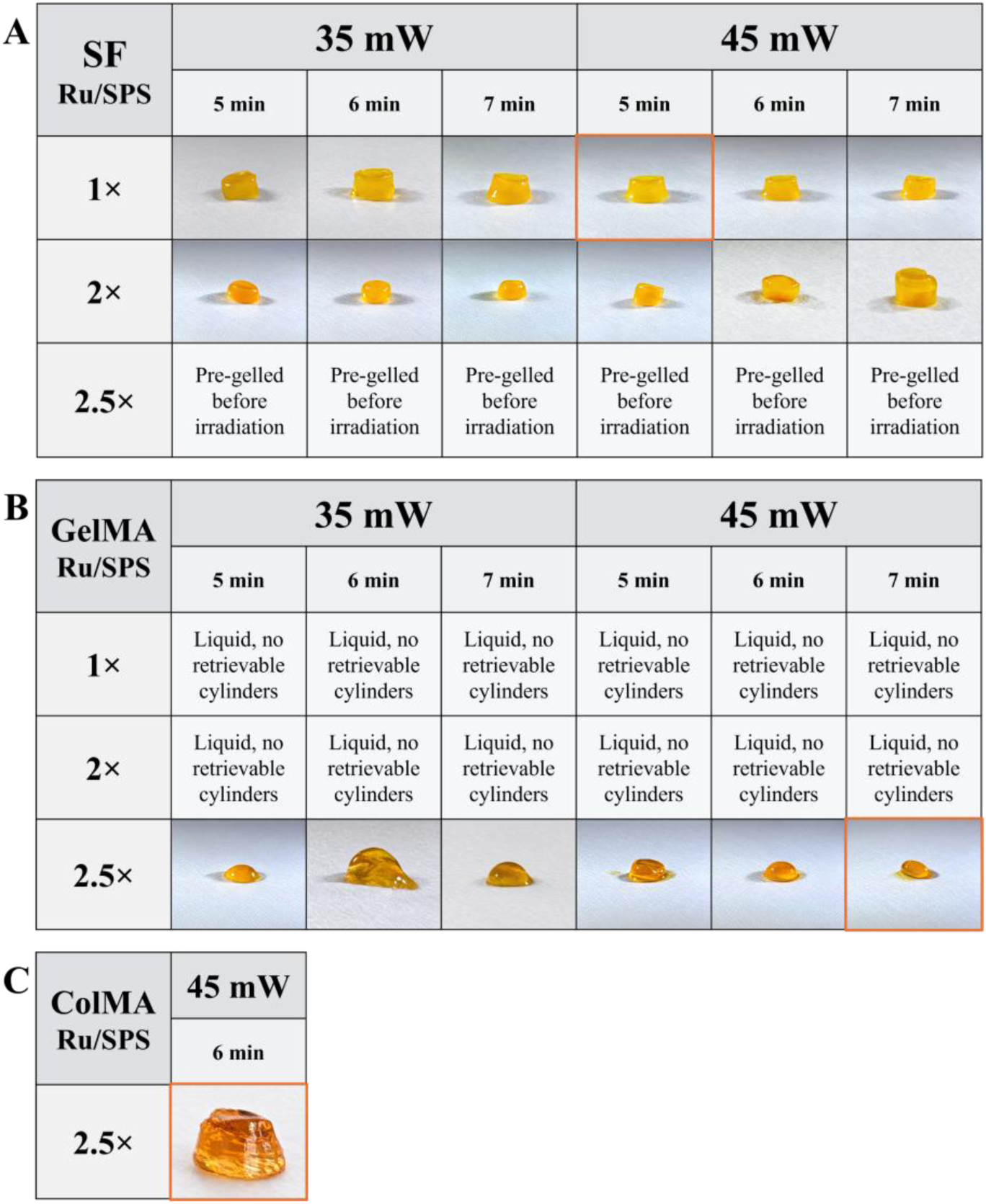
Screening and selection of visible-light photocrosslinking conditions for silk fibroin (A), GelMA (B), and ColMA (C). Cylindrical formulations were exposed to 430-nm light while varying Ru/SPS concentration, optical power (35 or 45 mW), and irradiation time (5–7 min). Photocrosslinking was considered successful when an intact, selfsupporting hydrogel could be recovered from the mold and retained its cylindrical geometry during handling. For SF, both 1× and 2× Ru/SPS generated retrievable constructs, whereas the 2.5× formulation pre-gelled before irradiation; 1× Ru/SPS, 45 mW, and 5 min was selected for subsequent characterization. GelMA formulations containing 1× or 2× Ru/SPS remained liquid under the tested conditions, whereas 2.5× Ru/SPS produced recoverable hydrogels; 45 mW for 7 min was selected. ColMA was evaluated at 2.5× Ru/SPS, 45 mW, and 6 min following the selected working formulation and was not subjected to a complete optimization matrix. Orange outlines indicate the conditions selected for downstream characterization.

**Figure 3.**
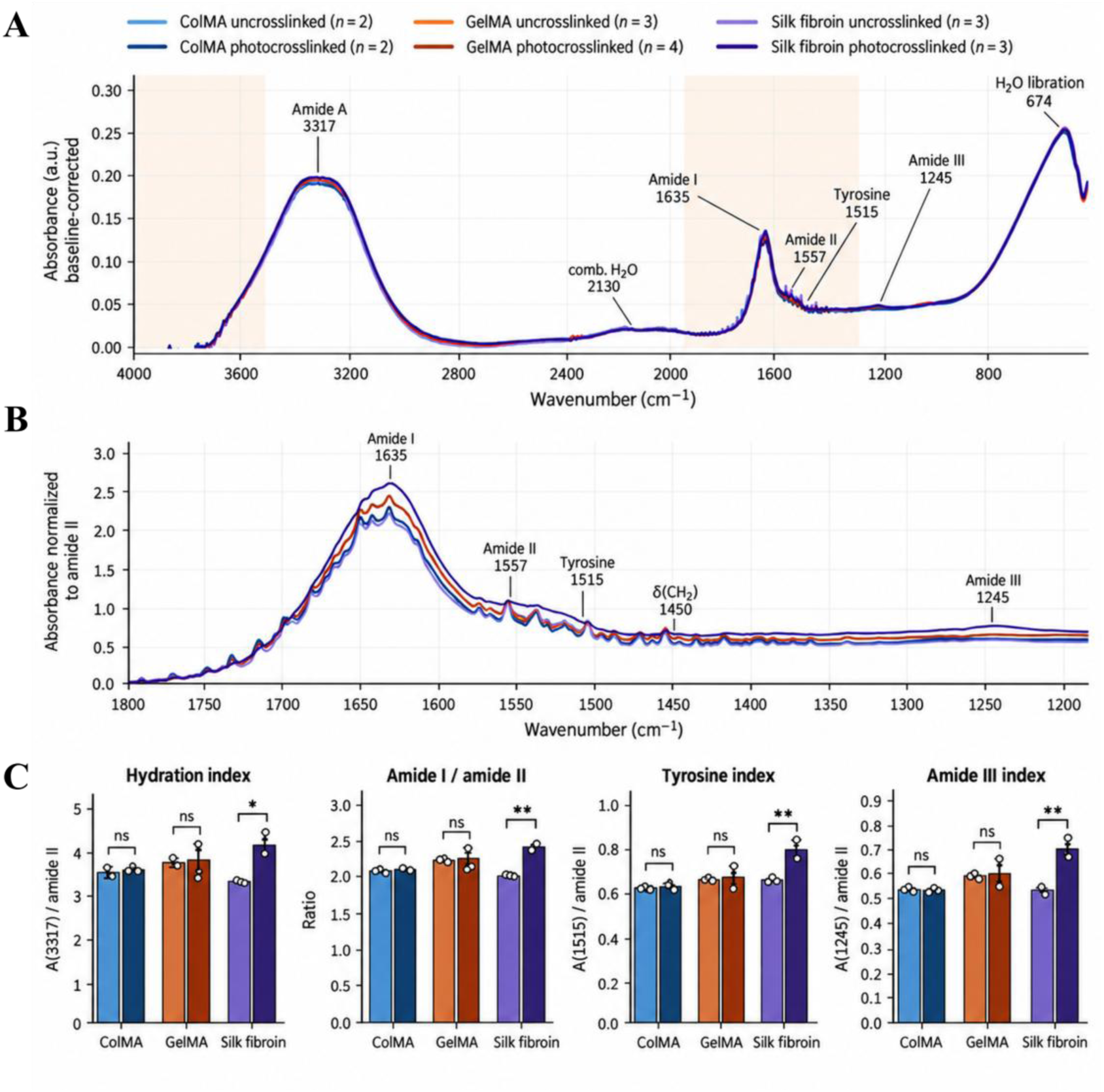
ATR-FTIR characterization of uncrosslinked and Ru/SPS-photocrosslinked protein formulations. (A) Baseline-corrected spectra of ColMA, GelMA, and silk fibroin (SF) are shown over the complete measured spectral range together with the protein-associated 1800–1200 cm⁻¹ region normalized to the Amide II band (B). (C) Characteristic bands include Amide A (∼3317 cm⁻¹), Amide I (∼1635 cm⁻¹), Amide II (∼1557 cm⁻¹), tyrosine (∼1515 cm⁻¹), Amide III (∼1245 cm⁻¹), and water-associated features. Quantitative spectral descriptors comprise the hydration index [A(3317)/Amide II], Amide I/Amide II ratio, tyrosine index [A(1515)/Amide II], and Amide III index [A(1245)/Amide II]. Individual data points and group means are shown. Sample sizes were ColMA, n = 2 uncrosslinked and n = 2 photocrosslinked; GelMA, n = 3 uncrosslinked and n = 4 photocrosslinked; and SF, n = 3 uncrosslinked and n = 3 photocrosslinked. Comparisons between uncrosslinked and photocrosslinked states within each material were performed using two-sided Welch’s t-tests. ns, not significant; p < 0.05; *p < 0.01.

**Figure 4.**
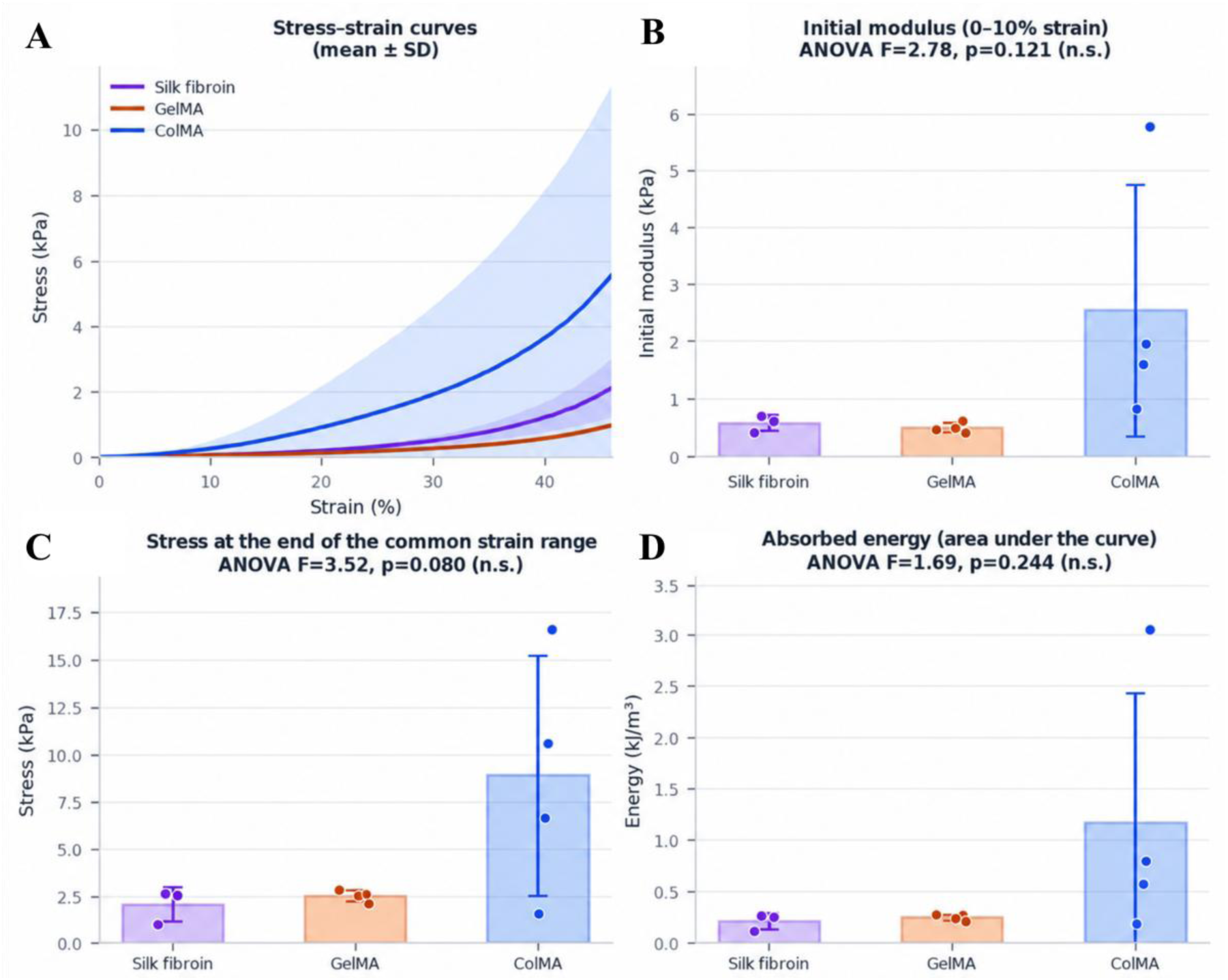
Compressive mechanical characterization of photocrosslinked protein hydrogels. (A) Engineering compressive stress–strain curves for silk fibroin (SF), GelMA, and ColMA, presented as mean ± SD. (B) Initial compressive modulus calculated by linear regression over the 0–10% strain interval. (C) Compressive stress measured at 45% strain. (D) Energy absorbed during compression, calculated as the area under the stress–strain curve over the common 0–45% strain interval. Bars represent mean ± SD and individual points represent independent hydrogel specimens (SF, n = 3; GelMA, n = 4; ColMA, n = 4). Comparisons among materials were performed using ordinary oneway ANOVA. No statistically significant between-material differences were detected for the three derived mechanical parameters (initial modulus: F = 2.78, p = 0.121; stress at 45% strain: F = 3.52, p = 0.080; absorbed energy: F = 1.69, p = 0.244). n.s., not significant.

**Figure 5.**
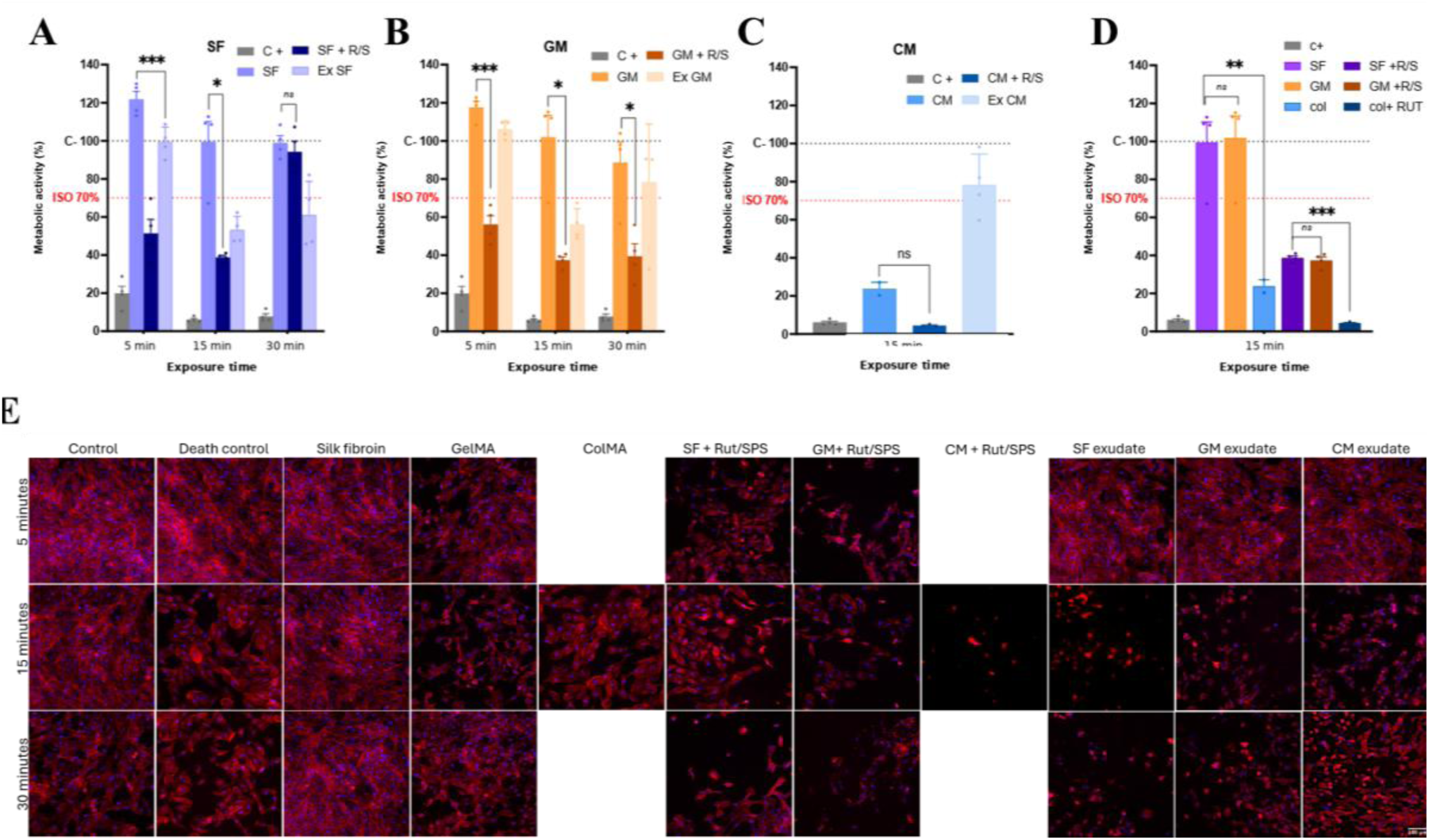
Short-term metabolic activity and cellular morphology of ARPE-19 cells exposed to the protein formulations. (A-C) Relative XTT metabolic activity following exposure to silk fibroin (SF), GelMA (GM), or ColMA (CM) formulations without photoinitiator, the corresponding unreacted Ru/SPS-containing precursor (+R/S), or medium conditioned by the photocrosslinked hydrogel (Ex). SF and GelMA were evaluated after 5, 15, and 30 min exposure; owing to limited material availability, the liquid ColMA precursor conditions were evaluated at 15 min. (D) Crossmaterial comparison at the matched 15-min exposure time. Complete culture medium was used as the negative/reference control and normalized to 100%, whereas 50% DMSO was used as the cytotoxic positive control (C+). The red dashed line indicates the 70% reference level derived from ISO 10993-5 and is shown only as an indicative benchmark; the assay was not conducted as a formal ISO 10993-5 cytotoxicity test. Bars represent mean ± SEM and points represent individual wells. Statistical analyses were performed as described in Materials and Methods. ns, not significant; p < 0.05; *p < 0.01; **p < 0.001. (E) Representative fluorescence images of ARPE-19 morphology following the corresponding exposures. F-actin was stained with Cy3-phalloidin (red) and nuclei with DAPI (blue). Scale bar, 100 µm.

**Figure 6.**
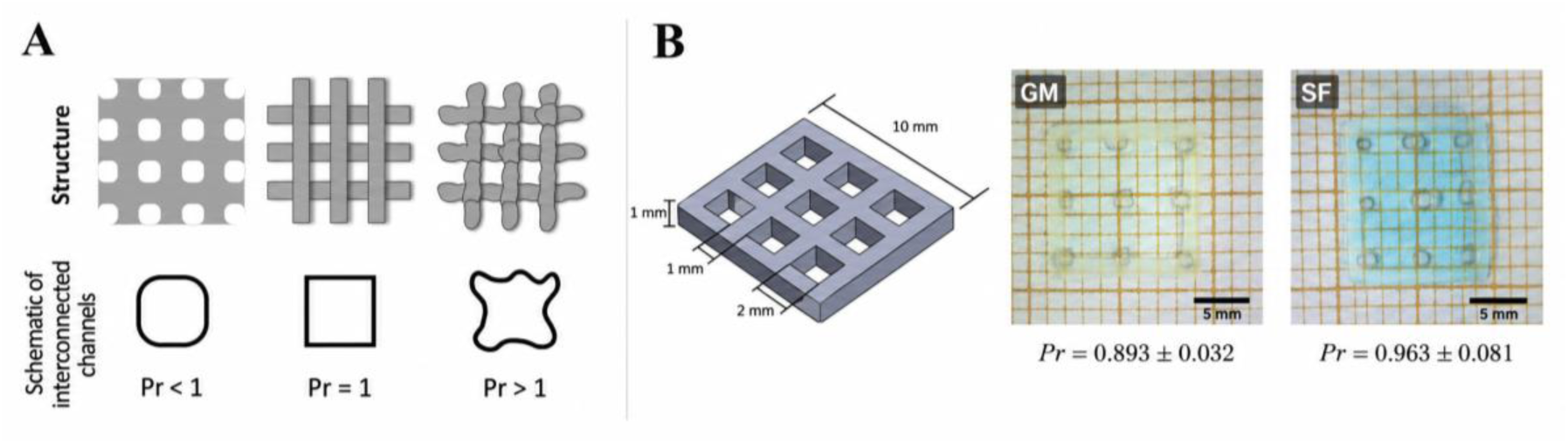
Printability assessment of GelMA and silk fibroin using a standardized 3 × 3 lattice geometry. (A) Schematic representation of the relationship between pore geometry and the printability index (Pr). A perfect square pore corresponds to Pr = 1; Pr < 1 indicates increasing pore rounding associated with material spreading, whereas Pr > 1 indicates increasingly irregular pore geometry. (B) Nominal lattice geometry and representative printed GelMA (GM) and silk fibroin (SF) constructs. Mean Pr values were 0.893 ± 0.032 for GelMA and 0.963 ± 0.081 for SF. Data were obtained from three independent printing experiments per material (n = 3 independent lattices), with nine pores analyzed as technical measurements within each lattice. Values are reported as mean ± SD. Scale bars, 5 mm.

**Figure 7.**
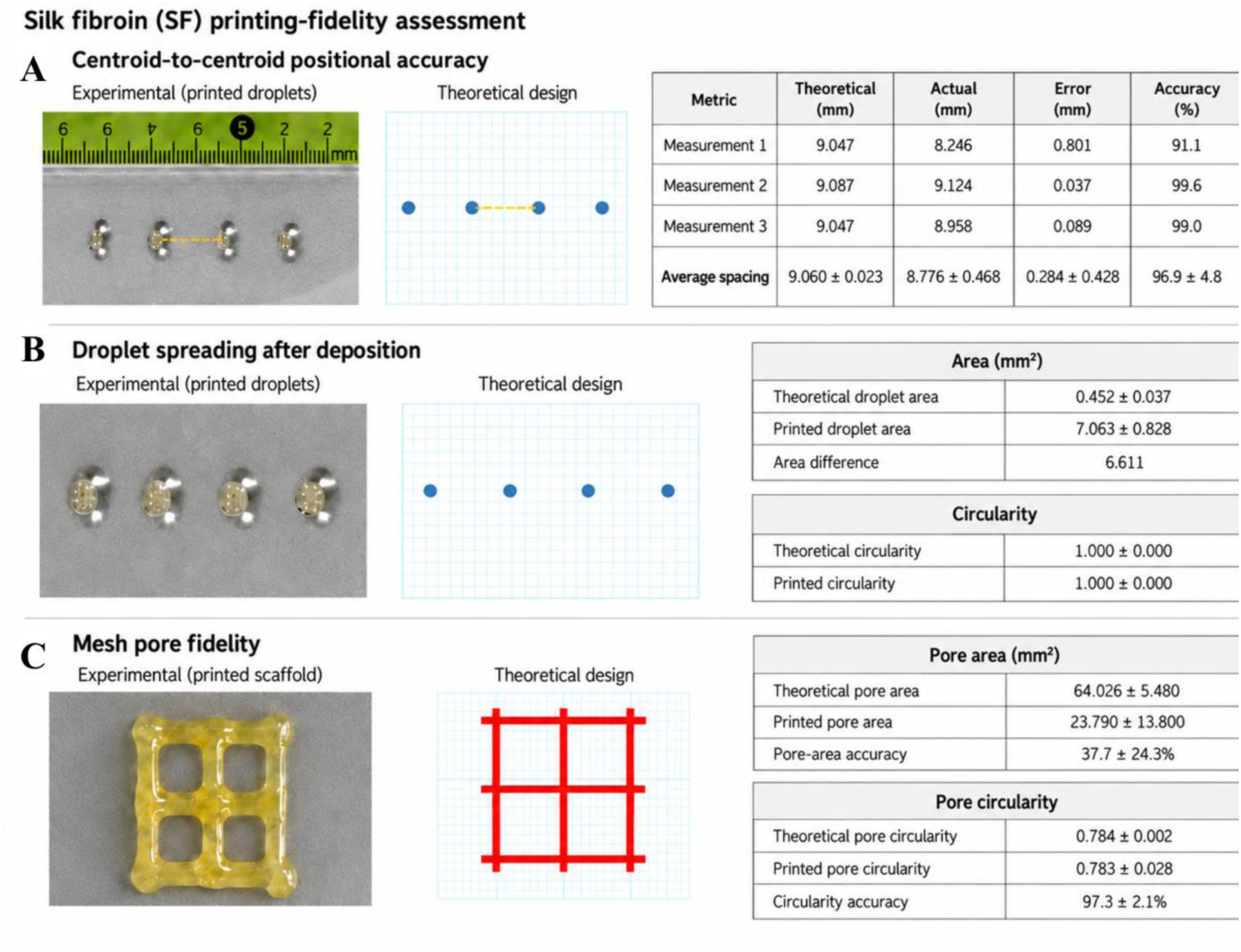
Quantitative assessment of positional and geometric printing fidelity in silk fibroin. (A) Centroid-to-centroid positional accuracy of a droplet-based pattern. Experimental droplet positions were compared with the corresponding theoretical CAD design; three adjacent centroid distances were analyzed, yielding an overall positional accuracy of 96.9 ± 4.8%. (B) Post-deposition droplet spreading. Theoretical and printed projected areas were compared together with circularity; printed droplets showed substantial area enlargement while retaining a circularity of 1.000 ± 0.000. Four droplets were analyzed. (C) Mesh pore fidelity. Experimental pore area and circularity were compared with the theoretical design. Pore-area accuracy was 37.7 ± 24.3%, whereas circularity accuracy was 97.3 ± 2.1%. Four pores were analyzed. Measurements within each pattern represent geometric technical measurements rather than independent printing replicates and were therefore analyzed descriptively.

Because the methacryloyl-associated C=C vibration overlaps the broad Amide I region in GelMA and ColMA, disappearance of a discrete C=C band was not used as standalone evidence of methacryloyl conversion.

### 2.7. Mechanical characterization

Mechanical behavior of the photocrosslinked hydrogels was evaluated by unconfined uniaxial compression using an Instron 4411 universal testing machine (Instron). Because the forces generated by the soft hydrogels were below the optimal working range of the conventional load-cell configuration, an analytical balance with 0.1-mg resolution and 250-g capacity (VWR) was positioned beneath the compression platen. The balance was placed on a granite support to minimize environmental vibration.

Specimens were prepared from 200 µL of the selected formulations using the cylindrical molds described above and photocrosslinked within the molds one day before mechanical testing. Samples were stored at 4 °C in PBS between photocrosslinking and testing. Compression was performed at a constant displacement rate of 1 mm/min, and force-related measurements were acquired at 1 Hz.

Balance readings were converted to compressive force according to:

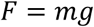

where *m* is the measured mass and *g* is gravitational acceleration. Engineering stress was calculated as:

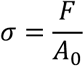

where the initial cross-sectional area was:

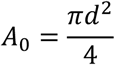

and engineering strain was calculated as:

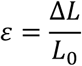

where *L*_0_ is the initial specimen height and Δ*L* is the applied displacement.

The initial compressive modulus was calculated as the slope of the linear regression fitted over the 0– 10% strain interval. For between-material comparisons at larger deformation, all stress–strain curves were analyzed over a common 0–45% strain interval, which was represented by all replicates of the three formulations. Compressive stress at 45% strain was extracted as a second mechanical descriptor. The energy absorbed during compression was calculated as the area under the stress-strain curve:

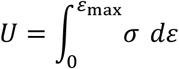

and expressed as energy per unit volume (kJ/m³).

### 2.8. ARPE-19 cell culture and short-term cytocompatibility assessment

#### 2.8.1. Cell culture

Human ARPE-19 retinal pigment epithelial cells (Cytion, Cat. No. 305025) were used as a reproducible adherent epithelial model for cytocompatibility assessment. The cells were not subjected to prolonged differentiation before the experiments and were therefore used as a cytocompatibility model rather than as a fully mature functional retinal pigment epithelium model.

Cells were maintained in DMEM/F12 (Gibco, Cat. No. 11320-033) supplemented with 10% FBS and 1% penicillin/streptomycin in a humidified incubator at 37 °C and 5% CO₂. Cells were passaged using trypsin (Gibco, Cat. No. 25200056).

For cytocompatibility experiments, ARPE-19 cells were seeded into 48-well tissue-culture plates (Thermo Scientific, Cat. No. 130187) at 30,000 cells per well in 300 µL complete medium and allowed to attach overnight. Independent plates were used for each exposure time.

#### 2.8.2. Experimental exposure conditions

Three material states were evaluated for each biomaterial: (i) the protein formulation without Ru/SPS, (ii) the corresponding formulation containing Ru/SPS before visible-light photocrosslinking, and (iii) conditioned medium obtained from the photocrosslinked hydrogel. Complete culture medium served as reference control and 50% dimethyl sulfoxide (DMSO) served as the cytotoxicity control.

Hydrogel-conditioned medium was generated by incubating four photocrosslinked 200-µL cylinders of each formulation in 4 mL complete culture medium for 24 h. The conditioned medium was subsequently recovered, frozen until use, and sterile-filtered through 0.22-µm syringe filters.

Exposure periods of 5, 15, and 30 min were selected to represent short residence times potentially encountered by cells during bioink preparation, reservoir loading, extrusion, and photocrosslinking. Accordingly, the experiment was not designed as a formal ISO 10993-5 extract-cytotoxicity test. The 70% reference level described in ISO 10993-5 was used only as an indicative benchmark for interpretation of relative metabolic activity.

Four wells were evaluated for each condition and exposure time unless otherwise stated. The medium reference at 5 min contained n = 3 wells; photoinitiator-free ColMA at 15 min contained n = 2; and ColMA containing Ru/SPS at 15 min contained n = 3. Because of limited ColMA availability, the liquid ColMA precursor and ColMA + Ru/SPS conditions were evaluated only at the 15-min time point.

#### 2.8.3. XTT metabolic activity assay

Following overnight attachment, culture medium was removed and cells were exposed to 200 µL of the corresponding test condition for 5, 15, or 30 min. The exposure solution was subsequently removed and replaced with 200 µL of freshly prepared XTT working solution from the CyQUANT™ MTT and XTT Cell Viability Assays kit (Invitrogen, Thermo Fisher Scientific, Cat. No. V13154), prepared according to the manufacturer’s instructions.

Cells were incubated with XTT for 4 h at 37 °C. Absorbance was measured at 450 nm using 660 nm as the reference wavelength with a SPECTROstar Omega microplate reader (BMG LABTECH). Corrected absorbance was calculated as:

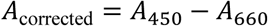

Relative metabolic activity was calculated against the culture-medium control from the corresponding exposure-time experiment:

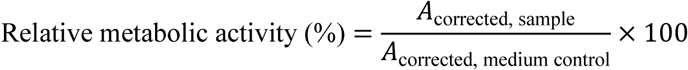

The XTT assay was interpreted as a measurement of relative metabolic activity and not as a direct cell-counting assay.

#### 2.8.4. Fluorescence assessment of cell morphology

Following completion of the XTT assay, the same cultures were fixed with 4% paraformaldehyde (PFA) for 20 min and washed three times with PBS. Cells were permeabilized with 0.1% Triton X-100 (Sigma-Aldrich, Cat. No. X100-5ML) for 30 min.

F-actin was stained using Cy3-conjugated phalloidin (Invitrogen, Cat. No. A34055) diluted 1:1000 in PBS for 90 min at room temperature protected from light. Nuclei were stained with DAPI (Thermo Scientific, Cat. No. 62248) at 1:1000 for 20 min. Fluorescence images were acquired using a Zeiss Colibri 7 equipped with an ORCA-Fusion camera (Hamamatsu Photonics).

These images were used as a qualitative orthogonal assessment of cell density, spreading, nuclear distribution, and F-actin organization. They were not treated as an independent quantitative cell-viability endpoint.

### 2.9. Lattice printability assessment

Printability of the selected GelMA and 2% (w/v) SF formulations was assessed using a standardized single-layer 3 × 3 lattice design based on established lattice-printability approaches [4,7]. The digital construct measured 10 × 10 mm, with a nominal height of 1 mm, filament width of 1 mm, and nine square pores measuring 2 × 2 mm.

The geometry was designed using Tinkercad (Autodesk), processed using EasyWare, and sliced using Cura. Nominal slicing parameters were: printing speed, 25 mm/s; travel speed, 25 mm/s; first-layer speed, 5 mm/s; infill speed, 20 mm/s; outer-wall speed, 20 mm/s; and inner-wall speed, 25 mm/s.

GelMA and SF were deposited through a sterile 23G blunt needle onto 22 × 50 mm glass coverslips. The extrusion system was operated at approximately 75 Hz and 15 V. A small quantity of blue dye was incorporated into the acellular formulations solely to improve contrast during macroscopic imaging.

Immediately after deposition, the thin lattices were photocrosslinked at 430 nm. GelMA constructs were irradiated for 3 min and SF constructs for 1 min. These shorter exposure times relative to the 200-µL bulk cylinders were selected because the printed structures contained substantially lower material volumes and thicknesses.

Three independent printing experiments were performed for each formulation (n = 3 independent lattices per material). Each lattice contained nine measurable pores. Individual pores within one construct were considered technical measurements, whereas the independently fabricated lattice was considered the experimental unit.

Printed constructs were photographed using a LINKMICRO LM210S magnification imaging system and analyzed using ImageJ (National Institutes of Health, USA; v. 1.48). The projected pore area *A*and perimeter *L*were measured, and the printability index was calculated according to Ouyang *et al.* [7]:

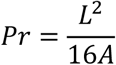

For a perfect square, *Pr* = 1. Values below 1 indicate increasing pore rounding associated with material spreading, whereas values above 1 indicate progressively irregular pore geometries. ColMA was not included in the lattice-printing experiment because insufficient material remained to generate a replicated printing series.

### 2.10. Independent assessment of printing positional and geometric fidelity

A separate SF printing experiment was analyzed to distinguish positional accuracy of the extrusion system from post-deposition deformation of the material. This dataset was generated during an earlier optimization stage using a different SF formulation and printing configuration and was therefore analyzed independently from the main GelMA/SF lattice experiment.

The formulation contained 2.5% (w/v) regenerated SF and was printed through a 23G needle at a nominal printing speed of 2 mm/s. The theoretical CAD designs and experimentally fabricated patterns were independently calibrated in ImageJ. Experimental images were calibrated using a physical ruler included in the acquired image, whereas theoretical Tinkercad images were calibrated using the digital design grid.

For the droplet-based pattern, centroid-to-centroid distances between adjacent elements were determined in both the theoretical design and the experimentally deposited pattern. Absolute positional error was calculated as:

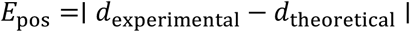

and positional accuracy as:

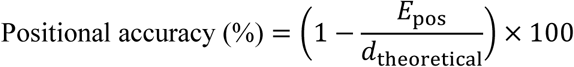

Projected droplet area was measured to quantify post-deposition spreading. Circularity was calculated as:

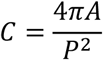

where *A*is projected area and *P*is perimeter; *C* = 1 represents a perfect circle.

The same analytical approach was applied to the mesh-based construct. Theoretical and experimental pore area and pore circularity were measured from calibrated images, and dimensional deviation and percentage agreement were calculated relative to the theoretical design. Three centroid-spacing measurements, four deposited droplets, and four mesh pores were analyzed. These values represented geometric measurements within the corresponding fabricated patterns rather than independent printing replicates and were therefore analyzed descriptively only.

### 2.11. Cell-laden bioprinting and spatial cell localization

Proof-of-concept cell-laden printing was performed using the selected GelMA and 2% (w/v) SF formulations prepared under aseptic conditions. Formulations were sterilized by filtration before cell incorporation.

ARPE-19 cells were harvested using trypsin, collected by centrifugation at 1200 rpm for 5 minutes, and labeled with Vybrant™ DiI Cell-Labeling Solution (Invitrogen, Thermo Fisher Scientific, Cat. No. V22885) according to the manufacturer’s protocol. Labeled cells were resuspended in the corresponding bioink at approximately 5.8 × 10⁵ cells/mL.

Approximately 400 µL of the cell-containing formulation was loaded into the printing reservoir and deposited through a sterile 23G blunt needle using the same 3 × 3 lattice geometry described above. Constructs were printed onto sterile glass coverslips and immediately photocrosslinked at 430 nm for 3 min for GelMA or 1 min for SF.

Immediately after fabrication, the constructs were examined using an Eclipse E200 fluorescence microscope (Nikon). DiI fluorescence was used to determine whether cells remained spatially associated with the deposited material and printed filament pattern. This assay was designed to assess successful cellular incorporation and spatial deposition and was not interpreted as a quantitative measurement of post-print cell viability. Cell-laden printing and DiI-based spatial localization were performed using both GelMA and SF formulations. Because the aim of this experiment was to provide a qualitative proof-of-concept of cell incorporation and spatial deposition rather than a quantitative comparison between bioinks, the SF construct is shown in Figure 8A as a representative example.

**Figure 8.**
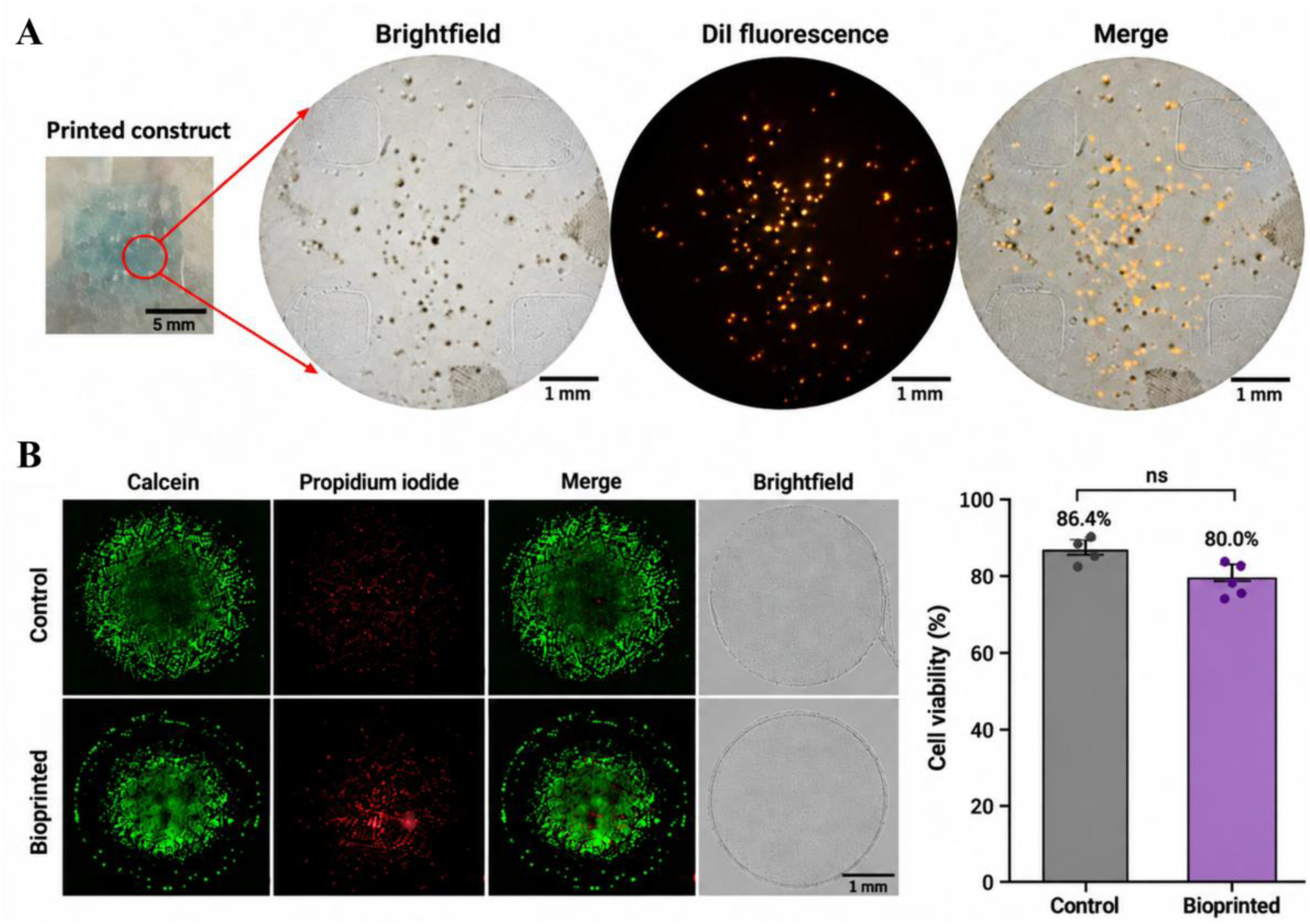
Spatial cell deposition and independent assessment of ARPE-19 viability following silk fibroin extrusion. (A) Representative silk fibroin proof-of-concept construct containing DiI-labeled ARPE-19 cells. Cell-laden printing and DiI-based spatial-localization experiments were performed using both SF and GelMA; SF is shown as the representative example. The printed construct, brightfield image, DiI fluorescence, and merged image demonstrate spatial association of the labeled cells with the deposited material. Scale bars, 5 mm for the macroscopic construct and 1 mm for microscopy images. (B) Live/Dead assessment of an independent sonication-gelled 2% SF experiment designed to isolate the effect of extrusion from Ru/SPS photocrosslinking. Constructs were deposited either manually through a needle (Control) or using the bioprinter (Bioprinted). Calcein-positive cells are shown in green and propidium-iodide-positive cells in red, together with merged and brightfield images. Mean viability was 86.4 ± 0.8% for manually deposited controls and 80.0 ± 3.2% for bioprinted constructs (n = 3 independent constructs per condition). Bars represent mean ± SD with individual replicate values. Groups were compared using a two-sided Welch’s t-test; ns, not significant. Scale bar, 1 mm.

**Figure 9.**
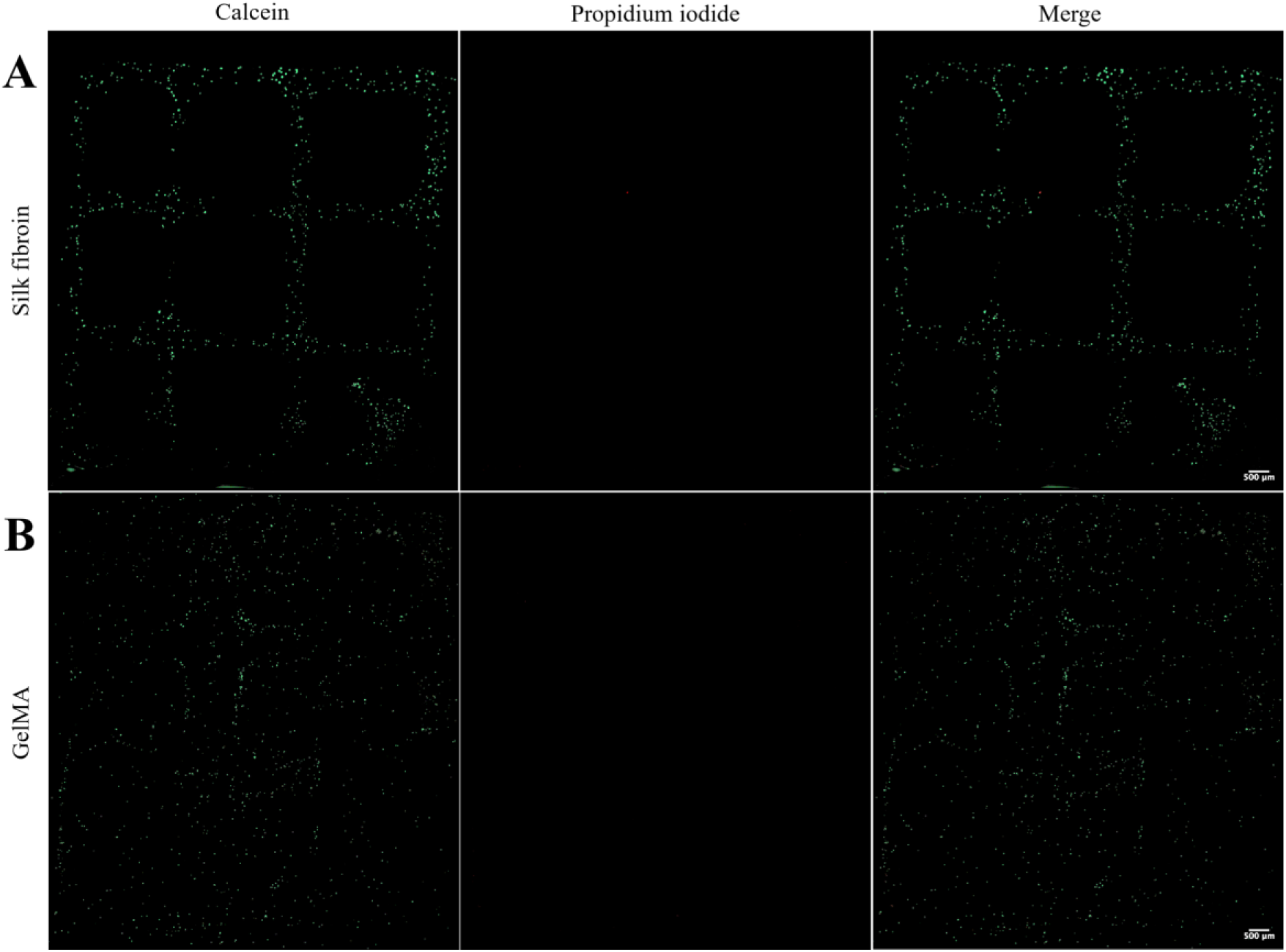
Same-day cell viability and spatial retention following the complete Ru/SPS-mediated biofabrication workflow. Representative fluorescence images of ARPE-19-containing silk fibroin (SF) (A) and GelMA (B) constructs approximately 30 min after extrusion and visible-light photocrosslinking. Cells were incorporated at 1 × 10⁶ cells/mL, printed through a 23G blunt needle, and photocrosslinked at 430 nm for 1 min for SF or 3 min for GelMA. Calcein fluorescence identifies viable cells, whereas propidium iodide (PI) identifies cells with compromised plasma membranes; merged images show only negligible PI signal in both formulations. SF retained a clearly recognizable grid-associated cellular distribution, whereas GelMA lost the original lattice organization after addition of culture medium. The experiment was interpreted qualitatively and was not subjected to inferential statistical analysis. Scale bars, 500 µm.

### 2.12. Independent validation of cell viability after extrusion

A separate experiment was performed to determine whether passage through the extrusion system itself caused a detectable reduction in ARPE-19 viability independently of Ru/SPS-mediated photocrosslinking. For this purpose, a 2% (w/v) SF formulation gelled by sonication was used rather than the photocrosslinked SF formulation employed in the main workflow.

SF was sonicated using an ultrasonicator (UCD-500, Biobase) under a 1-minute program with cycles of 2 seconds of active pulse (Pulse ON) and 5 seconds of pause (Pulse OFF), with an amplitude of 30% of the equipment’s maximum value. ARPE-19 cells were subsequently incorporated at 1 × 10⁶ cells/mL. The same cell-containing formulation was deposited either with the bioprinting system or manually through a needle, such that deposition method was the principal experimental variable.

Constructs were deposited onto P60 culture dishes, allowed to gel, covered with complete culture medium, and incubated overnight at 37 °C and 5% CO₂.

Cell viability was subsequently assessed using calcein-AM (Invitrogen, Cat. No. C3100MP) to label viable cells and propidium iodide (PI; Invitrogen, Cat. No. P3566) to identify cells with compromised plasma membranes. Samples were incubated with the staining solution following the manufacturer protocol (Invitrogen, Cat. No. L3224) protected from light. Fluorescence images were acquired using a Zeiss Colibri 7 equipped with an ORCA-Fusion camera (Hamamatsu Photonics).

Live and PI-positive cells were counted using ImageJ, and viability was calculated as:

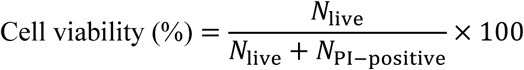

Three independent constructs were evaluated per condition (n = 3). Because this experiment used sonication-gelled SF rather than Ru/SPS-mediated photocrosslinking, it was interpreted specifically as an independent validation of the extrusion/deposition step and was not pooled with the main photocrosslinked cell-laden dataset.

### 2.13. Same-day viability assessment following the complete Ru/SPS biofabrication workflow

To assess cell status after integration of cell incorporation, extrusion, and Ru/SPS-mediated visible-light photocrosslinking, an additional qualitative Live/Dead experiment was performed using the final cell-containing SF and GelMA formulations.

ARPE-19 cells were incorporated into SF and GelMA at 1 × 10⁶ cells/mL and the formulations were printed through a sterile 23G blunt needle using the same single-layer 3 × 3 lattice workflow described above. Printed constructs were immediately photocrosslinked at 430 nm for 1 min for SF and 3 min for GelMA. Complete culture medium was subsequently added to the constructs.

On the same day as printing, at approximately 30 minutes, the constructs were stained with calcein-AM and PI. Calcein fluorescence was used to visualize metabolically active cells, whereas PI fluorescence was used to identify cells with compromised membranes. Samples were examined using a Zeiss Colibri 7 equipped with an ORCA-Fusion camera (Hamamatsu Photonics). Acquisition settings were maintained constant between the SF and GelMA constructs.

This experiment was used as a qualitative same-day viability and spatial-retention assessment of the complete photocrosslinked biofabrication workflow. No quantitative viability percentage was calculated from Figure 9; therefore, the images were not subjected to inferential statistical analysis.

### 2.14. Statistical analysis

Quantitative data are reported as mean ± standard deviation (SD) unless otherwise indicated. For the XTT experiments, Figure 5 presents individual observations together with mean ± standard error of the mean (SEM). Statistical analyses were performed using GraphPad Prism 8 (GraphPad Software) and Python 3 with SciPy. Statistical significance was defined as p < 0.05.

Given the small sample sizes and the unequal variances observed in several datasets, unequal-variance procedures were used where appropriate. Normality was not considered established solely on the basis of formal normality testing because of the limited diagnostic power associated with the small group sizes.

For ATR-FTIR analysis, each of the four normalized spectral indices was compared between the uncrosslinked and photocrosslinked conditions within each biomaterial using a two-sided Welch’s *t*-test.

For the mechanical-characterization dataset, initial compressive modulus, compressive stress at 45% strain, and absorbed energy over the 0–45% strain interval were compared among SF, GelMA, and ColMA using ordinary one-way ANOVA. This analysis corresponds to the statistical values reported in Figure 4.

For the ARPE-19 XTT dataset, statistical analysis was organized into predefined comparison families. Each experimental condition was first compared with the corresponding culture-medium control within the same exposure time using two-sided Welch’s *t*-tests followed by Benjamini-Hochberg correction for multiple comparisons. Within each biomaterial and exposure time, the formulation without photoinitiator, formulation containing Ru/SPS, and photocrosslinked-hydrogel-conditioned medium were compared using one-way Welch ANOVA followed, where appropriate, by pairwise Welch’s *t*-tests. Comparisons among different biomaterials within equivalent material states were performed using one-way Welch ANOVA followed by Games-Howell post hoc testing.

For the principal lattice-printability experiment, the independently printed lattice was considered the experimental unit (n = 3 per formulation), while the nine pores measured within each lattice were treated as technical measurements nested within that construct. Pore-level measurements were first summarized for each independent print and subsequently used to obtain construct-level descriptive values. Because the purpose of this experiment was to characterize printability rather than to establish material superiority with the small number of independent constructs, no inferential GelMA-versus-SF comparison was applied to the Pr values shown in Figure 6.

The independent SF positional-and geometric-fidelity dataset was analyzed descriptively because centroid distances, droplets, and pores represented multiple geometric measurements within individual fabricated patterns rather than independent printing replicates.

For the independent extrusion Live/Dead experiment shown in Figure 8, manual needle deposition and bioprinter deposition were compared using a two-sided Welch’s *t*-test, with three independent constructs per condition. The comparison was considered exploratory because of the small sample size. Figure 9 was analyzed qualitatively and was not subjected to inferential statistical testing.

## 3. Results

### 3.1. Material-specific selection of visible-light photocrosslinking conditions

Visible-light photocrosslinking was first screened to identify processing conditions capable of generating macroscopically stable hydrogels from silk fibroin (SF), gelatin methacryloyl (GelMA), and collagen methacryloyl (ColMA). Cylindrical samples were exposed to 430-nm light while varying Ru/SPS concentration, optical power, and irradiation time, and successful photocrosslinking was operationally defined by recovery of an intact, self-supporting cylinder that retained the imposed geometry during handling (Figure 2).

The three protein systems exhibited markedly different processing windows. SF produced recoverable constructs at relatively low photoinitiator concentrations. The 2.5× Ru/SPS formulation was already pre-gelled before irradiation and was therefore excluded from the irradiation screen, whereas both 1× and 2× formulations could be recovered after light exposure. Based on the screening matrix, 1× Ru/SPS, 45 mW, and 5 min was selected for subsequent bulk SF characterization.

GelMA required a substantially higher Ru/SPS concentration. Formulations containing 1× or 2× Ru/SPS remained liquid and did not yield retrievable cylinders under any of the tested combinations of irradiation power and time. At 2.5× Ru/SPS, self-supporting constructs were obtained, and 2.5× Ru/SPS, 45 mW, and 7 min was selected as the working condition. ColMA was evaluated under the higher-photoinitiator regime corresponding to the commercial formulation, and an intact hydrogel was obtained using 2.5× Ru/SPS, 45 mW, and 6 min. Because only this condition was evaluated for ColMA, it should be regarded as the selected working condition rather than as a fully optimized parameter set.

These results demonstrated that a common Ru/SPS photocrosslinking chemistry did not translate into a common processing condition across the three protein matrices. Instead, each material required a distinct combination of photoinitiator content and irradiation time to reach a macroscopically stable state. The final formulations carried forward to physicochemical, mechanical, and biological characterization are summarized in Table 1.

### 3.2 Visible-light photocrosslinking produces material-dependent ATR-FTIR changes

ATR-FTIR spectroscopy was used to compare the uncrosslinked and photocrosslinked states of ColMA, GelMA, and SF (Figure 3). The complete spectra displayed the characteristic absorption features of hydrated protein-based materials, including Amide A at approximately 3317 cm⁻¹, Amide I at 1635 cm⁻¹, Amide II at 1557 cm⁻¹, and Amide III at 1245 cm⁻¹. Additional features were observed at approximately 1515 cm⁻¹, associated with tyrosine residues, together with water-related bands around 2130 and 674 cm⁻¹.

Because absolute absorbance differed slightly among specimens, the 1800-1200 cm⁻¹ region was normalized to Amide II to facilitate comparison of protein-associated band shapes. Four spectral descriptors were subsequently evaluated: the hydration index, A(3317)/Amide II; the Amide I/Amide II ratio; the tyrosine index, A(1515)/Amide II; and the Amide III index, A(1245)/Amide II.

Neither ColMA nor GelMA showed statistically significant differences between the uncrosslinked and photocrosslinked states for any of these four spectral descriptors. In contrast, SF showed a consistent increase in all four indices following visible-light photocrosslinking. The hydration index increased significantly (*p* < 0.05), whereas the Amide I/Amide II ratio, tyrosine index, and Amide III index increased with higher statistical significance (*p* < 0.01; Figure 3).

Thus, within the analytical conditions used here, SF exhibited the clearest detectable spectral differences between the uncrosslinked and Ru/SPS-photocrosslinked states. In GelMA and ColMA, no statistically detectable change was resolved by the selected spectral indices. Importantly, the broad Amide I region overlaps the spectral region in which methacryloyl-associated C=C contributions are expected, preventing unambiguous assessment of methacryloyl conversion from these ATR-FTIR spectra alone.

### 3.3. The selected formulations formed compliant hydrogels with material-dependent mechanical variability

The compressive behavior of the photocrosslinked hydrogels was evaluated by unconfined uniaxial compression (Figure 4). All three materials exhibited nonlinear stress-strain responses, with stress progressively increasing as deformation increased. No abrupt macroscopic fracture event was observed within the analyzed strain ranges.

The initial compressive modulus, calculated over the 0-10% strain interval, was 0.60 ± 0.14 kPa for SF, 0.52 ± 0.09 kPa for GelMA, and 2.55 ± 2.20 kPa for ColMA. Although ColMA showed a numerically higher mean initial modulus, the difference among materials was not statistically significant (ANOVA F = 2.78, p = 0.121; Figure 4B).

A similar pattern was observed at larger deformations. At 45% strain, compressive stress reached 2.10 ± 0.89 kPa for SF, 2.57 ± 0.31 kPa for GelMA, and 8.88 ± 6.36 kPa for ColMA, without reaching statistical significance (F = 3.52, p = 0.080; Figure 4C). he corresponding energies absorbed over the 0–45% strain interval were 0.224 ± 0.079, 0.260 ± 0.028, and 1.155 ± 1.290 kJ/m³, respectively, and again no significant group effect was detected (F = 1.69, p = 0.244; Figure 4D).

The substantially larger dispersion observed for ColMA was evident across all three mechanical descriptors. One ColMA specimen reached an initial modulus of 5.78 kPa, terminal compressive stress of 16.65 kPa, and absorbed energy of 3.05 kJ/m³, whereas the remaining replicates were considerably lower. SF and GelMA showed substantially more closely clustered responses. Overall, the selected photocrosslinking conditions therefore generated soft, low-kPa hydrogels from all three protein systems, but reproducibility of the mechanical response differed markedly among formulations.

### 3.4. Cytocompatibility was strongly dependent on the presence of unreacted Ru/SPS

Short-term cytocompatibility was assessed in ARPE-19 cells to represent the interval during which cells could remain exposed to the formulations during preparation, loading, extrusion, and photocrosslinking (Figure 5). For each material, the precursor without photoinitiator, the corresponding formulation containing Ru/SPS before photocrosslinking, and conditioned medium obtained from the photocrosslinked hydrogel were evaluated. Relative metabolic activity was normalized to the matched culture-medium control, while 50% DMSO was used as a cytotoxic reference. The 70% level derived from ISO 10993-5 was used only as an orientative reference and not as a formal pass/fail criterion for this short-exposure assay.

The DMSO control reduced metabolic activity to 19.9 ± 3.7% at 5 min, 6.0 ± 0.7% at 15 min, and 7.7 ± 1.4% at 30 min, confirming that the assay detected a strong reduction in cellular metabolic activity. By contrast, photoinitiator-free SF remained close to or above the medium reference throughout the experiment, with values of 121.7 ± 4.2%, 99.4 ± 10.8%, and 98.8 ± 3.8% at 5, 15, and 30 min, respectively. Photoinitiator-free GelMA behaved similarly, reaching 117.6 ± 3.2%, 101.9 ± 11.5%, and 88.5 ± 10.8%, with no statistically significant difference from the corresponding medium control at any tested time point.

Addition of Ru/SPS markedly reduced the XTT signal. SF containing Ru/SPS decreased to 51.4 ± 7.3% at 5 min and 38.8 ± 1.1% at 15 min, both significantly below the corresponding photoinitiator-free SF formulation. At 30 min, metabolic activity reached 94.2 ± 5.5%, and the difference from photoinitiator-free SF was no longer statistically significant. GelMA containing Ru/SPS remained below the 70% reference at all three exposure times, with values of 56.0 ± 4.7%, 37.3 ± 1.8%, and 39.4 ± 6.6% at 5, 15, and 30 min, respectively. Each value was significantly lower than that of the corresponding photoinitiator-free GelMA formulation.

Because of limited ColMA availability, the liquid ColMA precursor conditions were evaluated only at 15 min. Photoinitiator-free ColMA produced 23.7 ± 3.4% relative metabolic activity, whereas ColMA containing Ru/SPS produced only 4.4 ± 0.4%. The within-ColMA comparison did not reach statistical significance because of the limited number of replicates, but the cross-material analysis showed a clear separation from SF and GelMA. At 15 min, SF and GelMA were statistically indistinguishable in both their photoinitiator-free states (99.4 ± 10.8% vs 101.9 ± 11.5%, p = 0.987) and their Ru/SPS-containing states (38.8 ± 1.1% vs 37.3 ± 1.8%, p = 0.788). ColMA was significantly lower than both materials without photoinitiator and after addition of Ru/SPS (Figure 5D).

Conditioned media from photocrosslinked hydrogels showed a different temporal response. At 5 min, metabolic activity remained near the medium control, reaching 99.3 ± 4.0% for SF, 105.8 ± 2.3% for GelMA, and 112.7 ± 2.0% for ColMA. Lower values were subsequently measured at longer exposure times: SF-conditioned medium reached 53.2 ± 3.6% and 61.0 ± 8.8% at 15 and 30 min; GelMA-conditioned medium reached 56.0 ± 4.2% and 78.2 ± 15.3%; and ColMA-conditioned medium reached 78.3 ± 8.1% and 44.4 ± 10.9%, respectively.

Fluorescence imaging of F-actin and nuclei was used as a qualitative morphological complement to the XTT assay (Figure 5E). Cells exposed to the photoinitiator-free SF and GelMA formulations generally retained an adherent morphology and extensive F-actin organization. Conditions containing Ru/SPS showed lower apparent cell densities and progressively altered morphology in several fields, particularly at longer exposure times. Because qualitative morphology did not fully parallel the metabolic data in every condition, these images were interpreted as an orthogonal morphological observation rather than as an independent quantitative viability endpoint. This analysis identified the uncrosslinked Ru/SPS-containing precursor state as a major short-term biological constraint within the tested processing workflow.

### 3.5. GelMA and silk fibroin could be printed as interconnected lattice structures

Following physicochemical, mechanical, and cytocompatibility screening, printing performance was evaluated for GelMA and SF using a common single-layer 3 × 3 lattice geometry (Figure 6). ColMA was not advanced to this stage because the available material was insufficient to generate a fully replicated printing dataset.

Both GelMA and 2% SF could be extruded into continuous interconnected structures in which the nominal pore pattern remained macroscopically identifiable. Quantification using the printability index, Pr, yielded 0.893 ± 0.032 for GelMA and 0.963 ± 0.081 for SF. Both values were below the theoretical value of 1 expected for an ideal square pore, indicating some degree of pore rounding following deposition. The SF value was numerically closer to unity, although both materials showed visible filament spreading at the macroscopic level. The analysis was therefore interpreted descriptively rather than as evidence of a statistically significant difference between materials.

Importantly, a Pr value approaching 1 described preservation of pore shape but did not establish preservation of absolute dimensions. This distinction motivated a separate quantitative analysis of positional and dimensional printing fidelity.

### 3.6. Silk-fibroin printing preserved positional accuracy despite substantial post-deposition spreading

An independent SF dataset was analyzed to separate errors in material placement from geometric changes occurring after deposition (Figure 7). In the droplet pattern, theoretical centroid-to-centroid distances of 9.047, 9.087, and 9.047 mm corresponded to experimentally measured distances of 8.246, 9.124, and 8.958 mm, respectively. The individual positional accuracies were therefore 91.1%, 99.6%, and 99.0%. Mean theoretical spacing was 9.060 ± 0.023 mm, compared with an experimentally measured spacing of 8.776 ± 0.468 mm, producing a mean absolute error of 0.284 ± 0.428 mm and an overall positional accuracy of 96.9 ± 4.8% (Figure 7A).

Despite this high positional agreement, pronounced dimensional changes occurred after deposition. The theoretical projected droplet area was 0.452 ± 0.037 mm², whereas the printed droplets occupied 7.063 ± 0.828 mm², corresponding to an average area increase of 6.611 mm². Circularity remained 1.000 ± 0.000 in both the theoretical and experimentally deposited droplets, showing that enlargement occurred without measurable loss of the approximately circular projected geometry (Figure 7B).

The effect of spreading became more evident in the mesh construct. The theoretical pore area was 64.026 ± 5.480 mm², whereas the experimentally measured pore area decreased to 23.790 ± 13.800 mm², corresponding to a pore-area accuracy of only 37.7 ± 24.3%. By contrast, pore circularity remained nearly unchanged, from 0.784 ± 0.002 in the theoretical design to 0.783 ± 0.028 after printing, corresponding to a circularity accuracy of 97.3 ± 2.1% (Figure 7C).

Taken together, these measurements showed that positional accuracy and final shape fidelity represented distinct components of printing performance. The dispensing system placed material close to the programmed coordinates, whereas substantial post-deposition spreading altered absolute dimensions and reduced open pore area. The principal source of geometric error in this experiment was therefore deformation after deposition rather than inaccurate spatial positioning. The measurements within each pattern represent geometric technical measurements and were not treated as independent biological or printing replicates.

### 3.7. Cell-laden printing preserved spatial cell deposition, while silk fibroin extrusion maintained short-term ARPE-19 viability

The capacity of the system to handle cell-containing formulations was next assessed using ARPE-19 cells. Cell-laden printing and DiI-based spatial localization were successfully performed with both SF and GelMA formulations. Figure 8A shows the SF construct as a representative proof-of-concept. DiI-labeled cells remained spatially associated with the deposited material, and the fluorescence signal reproduced the spatial distribution imposed by the printed construct, demonstrating that the cell-containing formulation could pass through the extrusion system while maintaining localization of the cellular component after deposition.

A separate experiment was then used to isolate the effect of the microfluidic process itself from Ru/SPS photocrosslinking. ARPE-19 cells were incorporated into 2% SF gelled by sonication and deposited either manually through a needle or using the bioprinter. Live/Dead imaging showed predominantly calcein-positive cells in both conditions, with a smaller PI-positive population (Figure 8B). The manually deposited control constructs showed cell viabilities of 85.5%, 86.9%, and 86.8%, corresponding to 86.4 ± 0.8%, whereas the bioprinted constructs showed 80.2%, 76.7%, and 83.0%, corresponding to 80.0 ± 3.2%. The difference was not statistically significant in the analysis represented in Figure 8.

These results indicated that, under the tested conditions, passage of the cell-containing SF formulation through the extrusion system did not produce a statistically detectable reduction in short-term ARPE-19 viability relative to manual deposition. Because this experiment used sonication-gelled SF rather than Ru/SPS-mediated photocrosslinking, it specifically isolated the extrusion/deposition component of the workflow rather than the complete photochemical process.

### 3.8. The complete Ru/SPS-mediated biofabrication workflow preserved cell viability but showed material-dependent structural retention

The final experiment evaluated cell-containing constructs after integration of the complete printing and visible-light photocrosslinking workflow (Figure 9). In both SF and GelMA constructs, fluorescence imaging showed extensive calcein-positive cell populations and only negligible propidium-iodide signal under the acquisition conditions. Thus, the cellular component remained predominantly viable after incorporation into the bioink, extrusion, and Ru/SPS-mediated visible-light stabilization.

The two materials nevertheless differed markedly in their ability to retain the printed spatial architecture. In SF, calcein-positive cells remained organized along the grid-like printed pattern, with the open regions of the lattice remaining clearly distinguishable. This spatial organization demonstrated that the printed SF construct maintained sufficient structural integrity after photocrosslinking and subsequent handling for the cellular distribution to remain associated with the original deposition path.

GelMA showed a contrasting response. Although the cells remained predominantly calcein positive, the original lattice architecture was not preserved after exposure to culture conditions. The GelMA construct lost structural integrity and dissolved/disassembled, resulting in redistribution of the cells away from the original filament pattern. Consequently, the absence of a retained grid in the GelMA fluorescence images reflected loss of scaffold architecture rather than an absence of viable cells.

Together, Figures 8 and 9 distinguish two separate biological requirements of the workflow. The independent sonicated-SF experiment showed that extrusion itself was compatible with high short-term cell viability, whereas the integrated Ru/SPS experiment showed that viable cell delivery alone was insufficient to guarantee successful biofabrication: post-crosslinking structural retention remained strongly material dependent.

### 3.9. Integrated definition of the biofabrication window

The sequential experimental workflow identified distinct material-and process-dependent constraints at different stages of biofabrication. Successful progression required not only formation of a stable photocrosslinked hydrogel, but also acceptable short-term cytocompatibility, extrusion through the printing system, preservation of geometric fidelity, and retention of the printed structure after fabrication. Although both SF and GelMA could be printed with viable cells, only SF maintained the grid-associated cellular architecture under the final tested conditions.

Together, these findings define the experimentally accessible biofabrication window as the set of conditions in which photocrosslinking, cytocompatibility, extrusion compatibility, geometric fidelity, and post-fabrication structural stability are simultaneously maintained.

## 4. Discussion

### 4.1. A biofabrication window emerges from coupled material and processing constraints

The present study investigated three protein-based formulations within a shared extrusion-biofabrication workflow to identify the constraints that determine whether a formulation can progress from liquid precursor to a stable, cell-containing printed construct. The principal finding is that successful biofabrication could not be attributed to a single property such as photocrosslinkability, stiffness, printability, or cytocompatibility. Instead, an experimentally accessible biofabrication window emerged from the interaction between formulation composition, visible-light stabilization, short-term biological exposure, extrusion behaviour, post-deposition dimensional stability, and preservation of cell viability.

This distinction is important because the three formulations were intentionally not treated as concentration-matched model polymers. SF, GelMA, and ColMA differed in polymer concentration, photoreactive chemistry, Ru/SPS concentration, and irradiation conditions. Consequently, the results should not be interpreted as an intrinsic ranking of the three protein materials. Rather, they show how experimentally relevant formulations encountered different limiting steps when subjected to the same sequential biofabrication framework. This approach reflects the practical reality of bioink development, in which a formulation is useful only if several requirements can be satisfied simultaneously [4–6].

The first constraint was already apparent during photocrosslinking. Despite using the same Ru/SPS initiating chemistry and 430-nm illumination, the three protein systems required different processing conditions to form self-supporting constructs. SF could be stabilized at the lowest Ru/SPS level tested for downstream characterization, whereas GelMA required the higher 2.5× condition and a longer irradiation period. ColMA also formed a recoverable hydrogel at 2.5× Ru/SPS, although this condition should be regarded as a working formulation rather than as an optimized processing window because material availability prevented a complete parameter screen.

The different behaviour is consistent with the distinct chemistry of these proteins. In GelMA and ColMA, Ru/SPS-generated radicals promote polymerization of methacryloyl groups, whereas in native SF the same photochemistry can oxidize tyrosine residues and generate covalent dityrosine crosslinks [15–18]. Network formation therefore depends not only on the amount of initiator and incident light but also on polymer concentration, accessibility and abundance of reactive groups, and the molecular organization of the precursor. The relatively low GelMA concentration used here, 16.67 mg/mL, is also substantially below the concentrations commonly used in many GelMA hydrogel and biofabrication studies [9]. A lower polymer content reduces the density of available network-forming groups and is therefore a plausible contributor to the longer exposure and higher Ru/SPS level required in the present formulation. Because the degree of methacryloyl substitution was not independently measured, however, its contribution cannot be separated from that of polymer concentration.

The SF response further illustrates why the useful processing space cannot be inferred from photocrosslinking chemistry alone. At excessive Ru/SPS concentration, the formulation had already undergone visible pre-gelation before irradiation. Thus, increasing initiator concentration did not simply extend the range of conditions capable of generating a hydrogel; beyond a certain point it reduced the usable handling window before printing. For an extrusion bioink, this distinction is critical because the precursor must remain sufficiently processable during loading and deposition while becoming stable rapidly once deposited.

### 4.2. Molecular and mechanical characterization supports network formation but also reveals analytical limitations

ATR-FTIR spectroscopy showed characteristic protein-associated bands in all three formulations, but the magnitude of the detectable spectral response differed among materials. No significant variation in the selected normalized indices was resolved for GelMA or ColMA, whereas SF showed higher hydration, Amide I/Amide II, tyrosine, and Amide III indices in the Ru/SPS-photocrosslinked state.

These observations should be interpreted conservatively. In GelMA and ColMA, the methacryloyl-associated C=C contribution lies within the broad Amide I region, where strong protein and water absorption limits direct assessment of double-bond conversion. Consequently, the absence of a significant change in the selected ATR-FTIR indices does not demonstrate absence of photocrosslinking. The transition from a liquid precursor to a manipulable hydrogel provides independent macroscopic evidence that a network was formed, but ATR-FTIR under the present hydrated conditions was not sufficiently specific to quantify methacryloyl conversion. Measurements of methacryloyl substitution before processing and more specific assessment of double-bond conversion would strengthen future characterization of GelMA and ColMA.

The SF spectral differences likewise should not be interpreted as direct quantification of dityrosine formation. The two analyzed states differed not only in irradiation history but also in the presence of Ru/SPS, and hydrated SF is particularly sensitive to differences in water content and conformational organization. Moreover, SF photocrosslinking can occur concurrently with slower self-assembly and secondary-structure rearrangement [17]. The increased normalized indices therefore demonstrate a measurable difference between the uncrosslinked and Ru/SPS-photocrosslinked SF states, but they do not independently identify the molecular origin of that difference. Direct dityrosine fluorescence, complementary structural analysis, or time-resolved assessment of SF secondary structure would provide a more specific description of the covalent and conformational contributions to network formation.

Mechanical testing confirmed that the selected conditions generated compliant hydrogels. The initial compressive moduli were in the low-kPa range, with SF and GelMA showing closely clustered values and ColMA displaying a higher mean accompanied by substantially greater inter-sample variability. Importantly, the absence of a statistically significant material effect should not be interpreted as evidence of mechanical equivalence. The relatively small sample numbers, particularly together with the large dispersion observed for ColMA, limit the ability to resolve differences between formulations.

The softness of the hydrogels is also consistent with their formulation context. GelMA was used at approximately 1.67% (w/v), substantially below many commonly reported GelMA formulations [9], while ColMA was used at only 3 mg/mL. In SF, mechanical properties can additionally evolve through processing-dependent self-assembly and maturation after covalent photocrosslinking [17,20]. Thus, the mechanical data are most useful here as characterization of the actual formulations used in the biofabrication workflow rather than as evidence of intrinsic mechanical differences among the three protein classes. Future studies should combine compression with rheological measurements of the precursor and post-crosslink network, because yield behaviour, shear thinning, recovery after extrusion, and time-dependent stabilization are more directly linked to printability than bulk compressive modulus alone [5,6].

### 4.3. The unreacted precursor state represents a critical biological constraint

The cytocompatibility experiments identified a second boundary of the biofabrication window. Photoinitiator-free SF and GelMA maintained metabolic activity close to that of the medium control over the short exposure intervals, whereas addition of Ru/SPS produced a marked decrease in XTT signal before photocrosslinking. This separation between polymer-only and Ru/SPS-containing conditions indicates that, for these two formulations, the unreacted initiating system was a major contributor to the acute biological response.

This result does not contradict previous reports demonstrating the suitability of visible-light Ru/SPS systems for cell-containing hydrogels [14–16]. Those applications generally minimize the interval during which cells remain exposed to freely diffusible initiating components by rapidly converting the precursor into a crosslinked network. The present XTT experiment deliberately evaluated the opposite situation: direct exposure of an adherent monolayer to the unreacted formulation during a short interval intended to represent preparation, loading, and pre-irradiation residence within the printing system. The data therefore identify time before stabilization as an important process variable. A formulation may support viable cells after photocrosslinking while still imposing a substantial biological burden if cells remain for prolonged periods in the unreacted precursor.

The SF time course deserves particular caution. Metabolic activity was markedly reduced at 5 and 15 min but approached the photoinitiator-free condition at 30 min. Because independent plates were used for the different exposure times, this pattern does not represent recovery of the same cellular population over time and should not be interpreted as cellular adaptation to Ru/SPS. Rather, it highlights the variability that can occur in short acute-exposure assays and reinforces the need to evaluate the biofabrication process as an integrated sequence rather than extrapolating from a single time point.

ColMA behaved differently because even the photoinitiator-free formulation produced a marked reduction in metabolic activity. This indicates that its biological response cannot be attributed to Ru/SPS alone. The low polymer concentration, acidic solubilization step, subsequent neutralization, ionic composition, and residual formulation components may all influence the acute response. Because ColMA material availability restricted both replication and the number of conditions evaluated, the present data identify a formulation-level compatibility problem but do not resolve its underlying mechanism.

Conditioned media from photocrosslinked hydrogels provided additional evidence that post-crosslinking exposure also requires consideration. Although the initial response was close to the medium control, reduced metabolic activity was observed for several extracts at longer cell-exposure intervals. Residual freely diffusible initiator, persulfate-derived species, unreacted formulation components, or other leachable products may contribute to this effect. Direct chemical quantification was outside the scope of this study, but the observation supports the inclusion of a post-crosslink washing step and quantitative assessment of residual Ru/SPS-derived species in future formulations.

The 70% reference shown in Figure 5 must therefore remain contextual rather than regulatory. The exposure design used here intentionally differs from the extraction procedure specified by ISO 10993-5 [22], and the experiment should not be described as demonstrating formal compliance or non-compliance with that standard. Similarly, ARPE-19 cells were used as a reproducible epithelial cytocompatibility model rather than as a mature retinal pigment epithelium. Their phenotype is strongly dependent on differentiation and culture conditions [21], so the present biological conclusions relate to acute material/process compatibility rather than retinal functional performance.

### 4.4. Printability cannot be represented by a single shape descriptor

Both GelMA and SF generated recognizable interconnected single-layer lattices, and their mean printability indices were close to the theoretical value of one for a square pore. Considered alone, these values would suggest relatively good preservation of the programmed pore geometry. The independent SF fidelity analysis, however, demonstrates why a single printability parameter is insufficient to characterize biofabrication performance.

In that experiment, centroid-to-centroid placement was highly accurate, yet deposited droplets occupied a much larger area than the corresponding theoretical elements. Likewise, pore circularity in the printed mesh remained close to its theoretical value while absolute pore area was markedly reduced. Shape and size therefore diverged: an element could retain essentially the expected circularity or pore geometry while undergoing substantial isotropic or near-isotropic expansion.

This finding is consistent with the broader view that bioink printability is multidimensional [4–7]. The Pr metric is useful because it provides a simple measure of deviation from square pore geometry, but it is fundamentally a shape descriptor derived from area and perimeter. It does not, by itself, establish dimensional accuracy, filament width, positional accuracy, pore size preservation, or structural stability after exposure to medium. A bioink can therefore produce a Pr value close to one while still deviating substantially from the CAD dimensions.

The SF geometric-fidelity dataset was obtained during an earlier optimization stage using a different SF concentration and printing configuration and should not be quantitatively extrapolated to the final 2% SF condition. Its value within this study is mechanistic: it separates positioning error from material deformation and demonstrates that the printer can place material near the programmed coordinates even when the material subsequently spreads. This distinction helps identify the appropriate target for future optimization. Improving stage positioning alone would not correct the observed dimensional error; the dominant opportunity lies in controlling precursor rheology and reducing the interval between deposition and network stabilization.

For this reason, future characterization of the system should combine construct-level replication with filament-width analysis, pore dimensions, layer height, dimensional accuracy relative to CAD, and rheological measurements before and after extrusion. This would provide a more complete process map than Pr alone and would better support extension from single-layer lattices toward multilayer constructs.

### 4.5. Extrusion and photocrosslinking impose distinct biological and structural requirements

The cell-containing experiments were deliberately separated into complementary stages to distinguish the effect of extrusion from that of the complete photochemical workflow. In the independent sonication-gelled SF experiment, manual deposition resulted in a mean viability of 86.4%, compared with 80.0% after bioprinter-mediated deposition. The difference was not statistically significant with three independent constructs per condition, suggesting that passage through the extrusion system was compatible with a high proportion of viable cells under the tested conditions.

Nevertheless, a non-significant result with this sample size should not be interpreted as proof of equivalence between the two deposition methods. The approximately six-percentage-point difference and the larger variability observed after bioprinting justify confirmation with greater experimental power. Extrusion-associated shear is a recognized determinant of cell injury, with nozzle geometry, material rheology, and flow conditions contributing jointly to the stress experienced by encapsulated cells [8]. The use of a 23G blunt needle in the final configuration is consistent with an effort to reduce flow resistance and recurrent obstruction, but shear stress was not directly measured here and therefore cannot be invoked as the demonstrated mechanism underlying the observed viability values.

The integrated Ru/SPS experiment adds a different level of evidence. Approximately 30 min after fabrication, both SF and GelMA constructs contained predominantly calcein-positive cells with only negligible PI signal. This qualitative observation indicates that cells could pass through formulation, extrusion, and visible-light photocrosslinking without widespread acute membrane damage. Importantly, this outcome differs from the marked metabolic reduction observed when adherent cells were exposed directly to unreacted Ru/SPS-containing precursor. The apparent discrepancy is informative: in the actual fabrication sequence, cells experience the unreacted state transiently, after which the construct is photocrosslinked and culture medium is added. Thus, the duration and physical context of initiator exposure are likely as important as nominal Ru/SPS concentration.

The same integrated experiment also demonstrated that biological survival and structural success are separate requirements. SF retained a recognizable grid-associated cellular distribution after fabrication, whereas GelMA lost the original lattice organization following addition of culture medium despite the presence of predominantly viable cells. The GelMA result therefore represents a structural rather than primarily biological failure of the workflow.

This observation should not be interpreted as evidence that SF is intrinsically more suitable for biofabrication than GelMA. The two formulations were not concentration matched and did not receive identical Ru/SPS doses or irradiation times. In particular, the GelMA concentration used here was low relative to many established GelMA bioinks [9], which likely contributed to insufficient network density and aqueous structural persistence. Optimization of GelMA polymer concentration, degree of methacrylation, irradiation conditions, and rapid layer-specific stabilization may therefore substantially shift its biofabrication window.

Conversely, the ability of SF to preserve the printed cellular architecture despite its low bulk modulus highlights the distinction between mechanical stiffness measured in compression and shape retention at the scale of a thin printed filament. A formulation does not necessarily require a high equilibrium modulus to maintain a single-layer geometry if network formation occurs rapidly enough to suppress lateral spreading and dissolution. Biofabrication performance must therefore be evaluated under the actual temporal and geometric conditions of printing rather than inferred solely from bulk material characterization.

### 4.6. Limitations and implications for future biofabrication studies

Several limitations define the scope of the conclusions. First, the study was not designed as a controlled head-to-head comparison of SF, GelMA, and ColMA. Polymer concentrations, reactive chemistries, Ru/SPS concentrations, irradiation durations, and formulation vehicles differed among materials. The results therefore describe the performance of specific experimentally accessible formulations and cannot be used to establish an intrinsic hierarchy among the underlying protein biomaterials.

Second, neither the degree of methacrylation of GelMA and ColMA nor the extent of photochemical conversion was quantified. This prevents direct correlation between reactive-group density, irradiation dose, and network properties. Future studies should incorporate characterization of functionalization together with photocrosslinking kinetics. For SF, direct assessment of dityrosine formation and time-dependent secondary-structure evolution would similarly help separate rapid covalent stabilization from subsequent conformational maturation [17].

Third, rheological characterization was not included in the present study. This is particularly relevant because viscosity, yield stress, shear thinning, and post-shear recovery are central determinants of extrusion behaviour and shape retention [5,6]. Combining rheology with the geometric metrics introduced here would enable the material-dependent biofabrication window to be mapped more quantitatively.

Fourth, ColMA availability restricted both statistical power and progression through the complete workflow. Its photocrosslinking condition was not comprehensively optimized, its cytocompatibility dataset was incomplete, and it was not evaluated in the replicated printing experiments. Conclusions regarding ColMA should therefore remain limited to the bulk formulation stages investigated here.

Finally, the biological experiments focused on acute or same-day responses. The Live/Dead data demonstrate short-term preservation of a predominantly viable cell population but do not establish long-term survival, proliferation, polarization, or retinal epithelial function. Future validation should include longer culture periods, quantitative viability at multiple post-printing time points, proliferation and morphology, and appropriate RPE phenotypic or barrier markers. Extension to multilayer constructs will additionally require stabilization during layer-by-layer deposition, because the present study focused on single-layer patterns.

These limitations also define a clear development path. Rather than optimizing polymer chemistry, microfluidic settings, and cell exposure independently, subsequent work should treat them as coupled variables. A useful formulation must remain extrudable during reservoir residence, stabilize rapidly enough to prevent spreading, achieve sufficient mechanical and aqueous stability after photocrosslinking, minimize exposure to freely diffusible initiating species, and preserve cellular viability. Systematic variation of these parameters, ideally supported by rheological and kinetic measurements, would enable construction of a quantitative processing map rather than a single optimized point.

Taken together, the present findings support the concept of a biofabrication window as an operational rather than purely material property. The window is bounded on one side by insufficient stabilization and geometric loss and on the other by excessive precursor reactivity, loss of processability, or biological exposure. Within those boundaries, extrusion accuracy, post-deposition fidelity, and cellular compatibility must all be satisfied simultaneously. The study therefore shows why successful extrusion bioprinting cannot be predicted from any isolated descriptor and provides an experimental framework for progressively defining workable conditions for visible-light photocrosslinkable protein bioinks.

## 5. Conclusions

This study demonstrates that the suitability of a protein formulation for extrusion bioprinting cannot be predicted from a single material property. Instead, a usable biofabrication window emerges from the combined requirements of precursor processability, visible-light photocrosslinking, short-term cytocompatibility, extrusion performance, geometric fidelity, and post-fabrication structural stability.

SF, GelMA, and ColMA required distinct Ru/SPS and irradiation conditions to form stable bulk hydrogels, confirming that a common photocrosslinking chemistry does not imply a common processing window. The unreacted Ru/SPS-containing state represented an important biological constraint, while printing experiments further showed that apparently favorable pore-shape metrics can coexist with substantial dimensional deformation. Cell-laden extrusion was compatible with high short-term ARPE-19 viability, but preservation of viable cells alone was insufficient to ensure successful construct formation: under the final tested conditions, SF retained the printed cellular architecture whereas GelMA lost its lattice organization after exposure to culture medium.

Overall, these findings support the concept of the biofabrication window as an operational property of the complete material–process–cell system rather than an intrinsic property of the biomaterial alone. The sequential framework developed here provides a practical basis for identifying the limiting steps of visible-light photocrosslinkable protein bioinks and for guiding their subsequent optimization toward more structurally stable and biologically compatible printed constructs.

## Author Contributions

Conceptualization: F.P. and G.A; methodology: J.M.-P. and S.M.-R.; investigation: J.M-P., S.M.-R., C.C.-D., A.S.-S, and A.P.-D.; resources: F.P.; writing—original draft preparation: J.M.-P.; writing— review and editing: S.M.-R., G.A. and F.P.; supervision: G.A. and F.P.; project administration: G.A. and F.P.; funding acquisition: G.A. and F.P. All authors have read and agreed to the published version of the manuscript.

## Data Availability Statement

Data will be available upon request.

## Acknowledgments

During the preparation of this manuscript/study, the authors used Scholar GPT 4 for the purposes of improving text syntax. The authors have reviewed and edited the output and take full responsibility for the content of this publication.

## References

1. Langer R, Vacanti JP. Tissue engineering. Science. 1993;260(5110):920–926.

2. Murphy SV, Atala A. 3D bioprinting of tissues and organs. Nat Biotechnol. 2014;32(8):773–785. doi:10.1038/nbt.2958.

3. Kang H-W, Lee SJ, Ko IK, Kengla C, Yoo JJ, Atala A. A 3D bioprinting system to produce human-scale tissue constructs with structural integrity. Nat Biotechnol. 2016;34(3):312–319. doi:10.1038/nbt.3413.

4. Gillispie G, Prim P, Copus J, Fisher J, Mikos AG, Yoo JJ, Atala A, Lee SJ. Assessment methodologies for extrusion-based bioink printability. Biofabrication. 2020;12(2):022003. doi:10.1088/1758-5090/ab6f0d.

5. Schwab A, Levato R, D’Este M, Piluso S, Eglin D, Malda J. Printability and shape fidelity of bioinks in 3D bioprinting. Chem Rev. 2020;120(19):11028–11055. doi:10.1021/acs.chemrev.0c00084.

6. Paxton N, Smolan W, Böck T, Melchels F, Groll J, Jungst T. Proposal to assess printability of bioinks for extrusion-based bioprinting and evaluation of rheological properties governing bioprintability. Biofabrication. 2017;9(4):044107. doi:10.1088/1758-5090/aa8dd8.

7. Ouyang L, Yao R, Zhao Y, Sun W. Effect of bioink properties on printability and cell viability for 3D bioplotting of embryonic stem cells. Biofabrication. 2016;8(3):035020. doi:10.1088/1758-5090/8/3/035020.

8. Blaeser A, Duarte Campos DF, Puster U, Richtering W, Stevens MM, Fischer H. Controlling shear stress in 3D bioprinting is a key factor to balance printing resolution and stem cell integrity. Adv Healthc Mater. 2016;5(3):326–333. doi:10.1002/adhm.201500677.

9. Nichol JW, Koshy ST, Bae H, Hwang CM, Yamanlar S, Khademhosseini A. Cell-laden microengineered gelatin methacrylate hydrogels. Biomaterials. 2010;31(21):5536–5544. doi:10.1016/j.biomaterials.2010.03.064.

10. Gaudet ID, Shreiber DI. Characterization of methacrylated type-I collagen as a dynamic, photoactive hydrogel. Biointerphases. 2012;7:25.

11. Cortella G, Lamparelli EP, Ciardulli MC, et al. ColMA-based bioprinted 3D scaffold allowed to study tenogenic events in human tendon stem cells. Bioeng Transl Med. 2025;10(1):e10723. doi:10.1002/btm2.10723.

12. Das S, Pati F, Choi Y-J, et al. Bioprintable, cell-laden silk fibroin-gelatin hydrogel supporting multilineage differentiation of stem cells for fabrication of three-dimensional tissue constructs. Acta Biomater. 2015;11:233–246. doi:10.1016/j.actbio.2014.09.023.

13. Zennifer A, et al. 3D bioprinting and photocrosslinking: emerging strategies & future perspectives. Biomater Adv. 2022;134:112576. doi:10.1016/j.msec.2021.112576.

14. Lim KS, Klotz BJ, Lindberg GCJ, et al. Visible light cross-linking of gelatin hydrogels offers an enhanced cell microenvironment with improved light penetration depth. Macromol Biosci. 2019;19(6):1900098. doi:10.1002/mabi.201900098.

15. Lim KS, et al. New visible-light photoinitiating system for improved print fidelity in gelatin-based bioinks. ACS Biomater Sci Eng. 2016;2(10):1752–1762. doi:10.1021/acsbiomaterials.6b00149.

16. Cui X, et al. Rapid photocrosslinking of silk hydrogels with high cell density and enhanced shape fidelity. Adv Healthc Mater. 2020;9(4):1901667. doi:10.1002/adhm.201901667.

17. Tran HA, Maraldo A, Ho TT-P, et al. Probing the interplay of protein self-assembly and covalent bond formation in photo-crosslinked silk fibroin hydrogels. Small. 2025;21(16):2407923. doi:10.1002/smll.202407923.

18. Steiner RC, Buchen JT, Phillips ER, Fellin CR, Yuan X, Jariwala SH. FRESH extrusion 3D printing of type-1 collagen hydrogels photocrosslinked using ruthenium. PLOS ONE. 2025;20(1):e0317350. doi:10.1371/journal.pone.0317350.

19. Pérez-Cortez J, Sánchez-Rodríguez V, Gallegos-Martínez S, Chuck-Hernández C, Rodriguez C, Álvarez M, Trujillo-de Santiago G, Vázquez-Lepe E, Martínez-López J. Low-cost light-based GelMA 3D bioprinting via retrofitting: manufacturability test and cell culture assessment. Micromachines. 2023;14(1):55. doi:10.3390/mi14010055.

20. Yonesi M, Garcia-Nieto M, Guinea GV, Panetsos F, Pérez-Rigueiro J, González-Nieto D. Silk fibroin: an ancient material for repairing the injured nervous system. Pharmaceutics. 2021;13(3):429.

21. Samuel W, Jaworski C, Postnikova OA, et al. Appropriately differentiated ARPE-19 cells regain phenotype and gene expression profiles similar to those of native RPE cells. Molecular Vision. 2017;23:60–89.

22. International Organization for Standardization. ISO 10993-5:2009. Biological evaluation of medical devices—Part 5: Tests for in vitro cytotoxicity.

